# Functional diversity of microboring *Ostreobium* algae isolated from corals

**DOI:** 10.1101/2020.09.18.303545

**Authors:** A. Massé, A. Tribollet, T. Meziane, M.L. Bourguet-Kondracki, C. Yéprémian, C. Sève, N. Thiney, A. Longeon, A. Couté, I. Domart-Coulon

**Affiliations:** Molécules de Communication et Adaptation des Microorganismes (MCAM), Muséum national d’Histoire naturelle (MNHN), CNRS (UMR7245); CP54, 63 Rue Buffon, 75005, Paris, France; IRD-Sorbonne Université (UPMC-CNRS-MNHN), Laboratoire IPSL-LOCEAN, 4 Place Jussieu, Tour 46-00, 5éme étage, 75005 Paris Cedex, France; Biologie des Organismes et Ecosystèmes Aquatiques (BOREA), Muséum national d’Histoire naturelle (MNHN), SU, UNICAEN, UA, CNRS (UMR7208), IRD; CP53, 61 rue Buffon, 75005, Paris, France

**Keywords:** *Ostreobium* sp., Ulvophyceae, coral skeleton, carbonate dissolution, carbon and nitrogen uptake, chlorophyll, fatty acids

## Abstract

The filamentous chlorophyte *Ostreobium* sp. dominates shallow marine carbonate microboring communities, and is one of the major agents of reef bioerosion. While its large genetic diversity has emerged, its physiology remains little known, with unexplored relationship between genotypes and phenotypes (endolithic *versus* free-living growth forms). Here, we isolated 9 strains affiliated to 2 lineages of *Ostreobium* (>8% sequence divergence of the plastid gene *rbc*L), one of which was assigned to the family Odoaceae, from the fast-growing coral host *Pocillopora acuta* Lamarck 1816. Free-living isolates maintained their bioerosive potential, colonizing pre-bleached coral carbonate skeletons. We compared phenotypes, highlighting shifts in pigment and fatty acid compositions, carbon to nitrogen ratios and stable isotope compositions (δ^13^C and δ^15^N). Our data show a pattern of higher chlorophyll *b* and lower arachidonic acid (20:4ω6) content in endolithic *versus* free-living *Ostreobium*. Photosynthetic carbon fixation and nitrate uptake, quantified via 8h pulse-labeling with ^13^C-bicarbonate and ^15^N-nitrate, showed lower isotopic enrichment in endolithic compared to free-living filaments. Our results highlight the functional plasticity of *Ostreobium* phenotypes. The isotope tracer approach opens the way to further study the biogeochemical cycling and trophic ecology of these cryptic algae at coral holobiont and reef scales.

## Introduction

The microscopic algal biodiversity dissolving actively (or eroding) shallow-water carbonates is a cryptic yet essential component of reef functioning, becoming dominant with the general decline of coral reefs worldwide (Leggat *et al*., 2019; Tribollet *et al*., 2019). Pioneer morphological observations showed that bioeroding filaments of the chlorophyte *Ostreobium* are ubiquitous inside the skeleton of tropical coral reef-builders, both in dead and actively growing colonies (Odum & Odum, 1955; Lukas, 1974; Le Campion-Alsumard *et al*., 1995a; reviewed in Tribollet, 2008; Golubic *et al*., 2019). Molecular data have recently accumulated, based on amplicon sequencing of plastid encoded gene markers (*rbc*L, *tuf*A, UPA and 16S rRNA), revealing *Ostreobium* ubiquity in the core microbiome of tropical corals and its high genetic diversity, delimiting an entire Ostreobineae suborder within the Bryopsidales in the class Ulvophyceae (Gutner-Hoch & Fine, 2011; Marcelino & Verbruggen, 2016; Sauvage *et al*., 2016; del Campo *et al*., 2017; Marcelino *et al*., 2017; Verbruggen *et al*., 2017; Gonzalez-Zapata *et al*., 2018; Marcelino *et al*., 2018; Massé *et al*., 2018). By chemical means, *Ostreobium* filaments actively penetrate reef carbonates ranging from limestone rocks to seashells and coral skeletons, creating galleries a few micrometers in diameter (Tribollet, 2008), thus living as true boring endoliths (i.e. euendoliths also called microborers; Golubic *et al*., 1981). Surprisingly, filaments of this photosynthetic chlorophyte can even be detected in microboring communities down to 200 m depth in tropical ecosystems (Littler *et al*., 1985; Vogel *et al*., 2000; reviewed in Tribollet *et al*., 2011). Occasionally, *Ostreobium* filaments can exit carbonate skeletons of coral holobionts or reef rubble to become epilithic (Kobluk & Risk, 1977) and free-living filaments can be detected in the environment in seawater or benthic biofilms (Massé *et al*., 2018). Filaments can also be released from their calcium carbonate substratum in culture (Kornman & Sahling, 1980; Sauvage *et al*., 2016). In declining reefs impacted by coral bleaching events and overfishing of algal grazers (Hughes *et al*., 2017; Roth *et al*., 2018), the prevalence of free-living *Ostreobium* filaments is likely to increase after detachment of epilithic filaments emerged from damaged coral colonies or reef rubble (Leggat *et al*., 2019), as a result of wave action during cyclone or storm events.

In massive adult coral colonies with a slow-growth, the endolithic layer dominated by bioeroding *Ostreobium* filaments forms visible green bands just beneath the coral tissues (Lukas, 1974; Le Campion *et al*., 1995a). In contrast, in fast-growing branching corals those green bands are absent, but *Ostreobium* filaments are still present, although at decreased abundance in the skeleton towards branch tips (Godinot *et al*., 2012; Massé *et al*., 2018). Recently, Massé *et al*. (2018) showed the horizontal transmission of *Ostreobium* from benthic biofilms or propagules dispersed in seawater to a fast-growing *Pocillopora* coral host. These microboring algae penetrate first into the skeleton of coral recruits, as soon as the primary polyp is formed after larval metamorphosis (Massé *et al*., 2018) and then follow the coral vertical extension in order to access enough light to survive (Halldal, 1968; Shibata & Haxo, 1969; Shashar & Stambler, 1992). Several functional roles have been suggested for *Ostreobium* (and other microborers) in living corals, such as nutrient recycling (Ferrer & Szmant, 1988) and a possible ectosymbiotic relationship between microboring communities dominated by *Ostreobium* and their coral host (Schlichter *et al*., 1995; Fine & Loya, 2002; Sangsawang *et al*., 2017). However, assimilation of inorganic carbon and nitrogen has not been quantified for individual, genetically referenced *Ostreobium* members of the skeleton microbiome, and sites of putative active transfer of metabolites from algal filament to host tissue are yet to be demonstrated in live reef corals. By contrast, in dead coral skeletons, the ecological roles of *Ostreobium* dominated assemblages have been more intensively studied. At complex community level, this microscopic alga is indeed, in synergy with bioeroding sponges and the grazing macrofauna such as parrotfishes, one of the major agents of bioerosion and calcium carbonate recycling (Tribollet & Golubic, 2005; Schönberg *et al*., 2017). Together with other microboring phototrophs, it is also an important benthic primary producer and thus, an important keystone in coral reef food web (Odum & Odum, 1955; Vooren, 1981; Tribollet *et al*., 2006; Clements *et al*., 2016).

To date, very little is known about the functional diversity of *Ostreobium* algae. Functional traits such as pigment and fatty acid compositions may vary in endolithic *versus* free-living growth habit and between specific genetic lineages. Moreover, it is not clear to which extent carbon (C) and nitrogen (N) sources and uptake rates may change to cover contrasting *Ostreobium* energy needs as endolithic filaments within carbonate substrate *versus* free-living filaments in seawater. Analyses of fatty acids and C and N stable isotopes could provide information on autotroph and/or heterotroph sources of carbon and nitrogen for *Ostreobium* filaments depending on their habitat. Pigment composition, and inorganic carbon and nitrogen uptake may also indicate adaptation of metabolic activity. Phenotypic studies of *Ostreobium* genetic lineages isolated from corals are thus crucial to better understand the role of these carbonate microboring algae at coral holobiont and reef ecosystem scales, and how they are impacted by environmental changes (Schönberg *et al*., 2017; Pernice *et al*., 2019; Ricci *et al*., 2019). This requires to compare in controlled laboratory settings the physiology of contrasting growth forms (phenotypes) of *Ostreobium*, i.e. endolithic filaments colonizing live coral colonies and reef carbonates *versus* free-living filaments.

Mono-algal cultures of *Ostreobium quekettii* Bornet and Flahaut 1889, which is the type species of the genus *Ostreobium*, were initially isolated from shells of a temperate mollusk in Brittany (France) (Bornet & Flahaut, 1889). Free-living strains 6.99 and B14.86 designated as *Ostreobium quekettii* were later used for further morphology and reproduction studies (Kornmann & Sahling, 1980), and trophic potential and photoecology investigations (Schlichter *et al*., 1997). Other free-living *Ostreobium* strains have since been isolated from reef-collected marine carbonates, genotyped with plastid encoded *tuf*A, UPA and 16S rRNA gene markers (Sauvage *et al*., 2016; Marcelino & Verbruggen, 2016) and then used for chloroplast genome sequencing and phylogenetic studies (Marcelino *et al*., 2016; del Campo *et al*., 2017; Verbruggen *et al*., 2017). Cultures of the endolithic form of *Ostreobium* have however seldom been characterized physiologically, except for a very recent study on strain 6.99 of *Ostreobium quekettii* Bornet and Flahaut 1889, that showed filament-driven processes of coral carbonate dissolution–reprecipitation and calcium transport (Krause *et al*., 2019).

In this study we developed an *in vitro* approach to compare the physiological characteristics of endolithic *versus* free-living *Ostreobium* filaments isolated from the fast growing, small polyp coral model species *Pocillopora acuta* Lamarck 1816. We genotyped nine strains of *Ostreobium* based on amplicon sequencing of the *rbc*L plastid gene marker, and characterized successive subcultures of these strains in either endolithic (coral carbonate eroding) or free-living growth habit to provide (see Table 1): (i) photosynthetic and accessory pigment composition, (ii) fatty acid composition, (iii) bulk tissue δ^13^C and δ^15^N stable isotope values and (iv) inorganic C and N assimilation patterns, measured via uptake of ^13^C-bicarbonate and ^15^N-nitrate stable isotope tracers.

**Table 1:** Analyses carried out per *Ostreobium* strain in endolithic *versus* free-living growth form. Controls are either bleached coral carbonate skeleton or residual skeletal organic matrix (after carbonate decalcification) of *Pocillopora acuta* host. (Strain 022 was lost since the analyses); subcult.: replicate subculture, techn. replicates: technical replicates.

| *Ostreobium*<br>strain code | Phylogenetic<br>affiliation<br>(*rbcL* gene #<br>Genbank) | MNHN RBCell<br>voucher # | Growth<br>form | Photosynthetic<br>and accessory<br>pigments | Fatty acids | Stable isotope values<br>($\delta^{13}\text{C}$ and $\delta^{15}\text{N}$) | <sup>13</sup>C -bicarbonate and<br><sup>15</sup>N-nitrate<br>assimilation | |
| --- | --- | --- | --- | --- | --- | --- | --- | --- |
|  |  |  |  |  |  |  | Light<br>condition | Dark<br>condition |
| 010 | Clade P1<br>(MK095212) | MNHN-ALCP-2019-873.3 | Free-living | + | + | + (2 subcult., each with<br>3 techn. replicates) | + (3 techn.<br>replicates) |  |
|  |  |  | Endolithic | + | + | + (2 subcult., each with<br>1-4 techn. replicates) | + |  |
| 017 | Clade P1<br>(MK095214) | MNHN-ALCP-2019-873.4 | Free-living |  | + |  |  |  |
|  |  |  | Endolithic |  | + |  |  |  |
| 018B | Clade P1<br>(MK095215) | MNHN-ALCP-2019-873.6 | Free-living | + (2 subcult.) | + (3 subcult.) | + |  |  |
|  |  |  | Endolithic | + | + (2 subcult.) |  |  |  |
| 019 | Clade P1<br>(MK095213) | MNHN-ALCP-2019-873.8 | Free-living | + (2 subcult.) | + | + |  |  |
|  |  |  | Endolithic | + | + |  |  |  |
| 05 | Clade P1<br>(MK095217) | MNHN-ALCP-2019-873.1 | Free-living | + | + (2 subcult.) | + (2 subcult.) | + | + |
|  |  |  | Endolithic |  | + | + (2 subcult.) | + | + |
| 018C | Clade P1<br>(MK095216) | MNHN-ALCP-2019-873.7 | Free-living | + (2 subcult.) |  |  |  |  |
|  |  |  | Endolithic |  | + |  |  |  |
| 022 | Clade P1<br>(MK095219) | NA | Free-living |  |  | + (2 subcult.) | + | + |
|  |  |  | Endolithic |  |  | + (2 subcult.) | + | + |
| 018A | Clade P12<br>(MK095218) | MNHN-ALCP-2019-873.5 | Free-living |  | + (3 techn. replicates) | + (2 subcult.) | + | + |
|  |  |  | Endolithic |  |  | + (2 subcult.) | + | + |
| 06 | Clade P14<br>(MK095220) | MNHN-ALCP-2019-873.2 | Free-living |  | + (2 subcult.) | + (2 subcult.) | + | + |
|  |  |  | Endolithic |  | + | + (2 subcult.) | + | + |
| Substrate controls for endolithic growth forms |  |  |  | Bleached coral<br>skeleton (n=3) | Bleached coral<br>skeleton (n=4) | Bleached coral skeleton (n=3)<br>Skeletal organic matrix (n=4) | Skeletal organic<br>matrix (n=3) |  |
| Corresponding data<br>(Figures, Tables and Supplementary information) |  |  |  | Fig. 2; Fig. S2;<br>Table S1 & S2 | Fig. 3; Fig. S3; Table<br>S3 | Fig. 4; Table S4& S5 | Fig. 5; Table S6 |  |

## Results

### Isolation and genotyping of *Ostreobium* strains to species-level *rbc*L clades

A total of nine *Ostreobium* strains were isolated from branch tips of *Pocillopora acuta* corals from long-term aquarium cultures (Aquarium Tropical, Palais de la Porte Dorée, Paris, Fr, called ATPD-aquarium). After removal of coral tissues, branched filaments with typical *Ostreobium* morphology emerged after ∼3 weeks from the skeleton of 2 out of 3 colonies (*Ostreobium* filaments did not emerge from the skeleton of one of the 3 colonies), forming yellowish-green tufts of filaments which were pulled out or cut with a scalpel to initiate cultures in free-living form. Siphoneous filaments had a diameter varying between 5 and 12 µm (Fig. 1b), with small disc-shaped chloroplasts visible in the periphery of siphons, against their inner sheath. Reproduction by spore formation was not observed. Mono-algal cultures, established via serial sub-culturing (successive passages) of such *Ostreobium* filaments, have been propagated *in vitro* in free-living and endolithic forms (see below) since September 2016, with vouchers deposited in the Museum national d’Histoire naturelle (Paris, France) RBCell collection of microalgae and cyanobacteria (MNHN-ALCP-2019-873.1 to MNHN-ALCP-2019-873.8; see Table 1).

**Figure 1:**
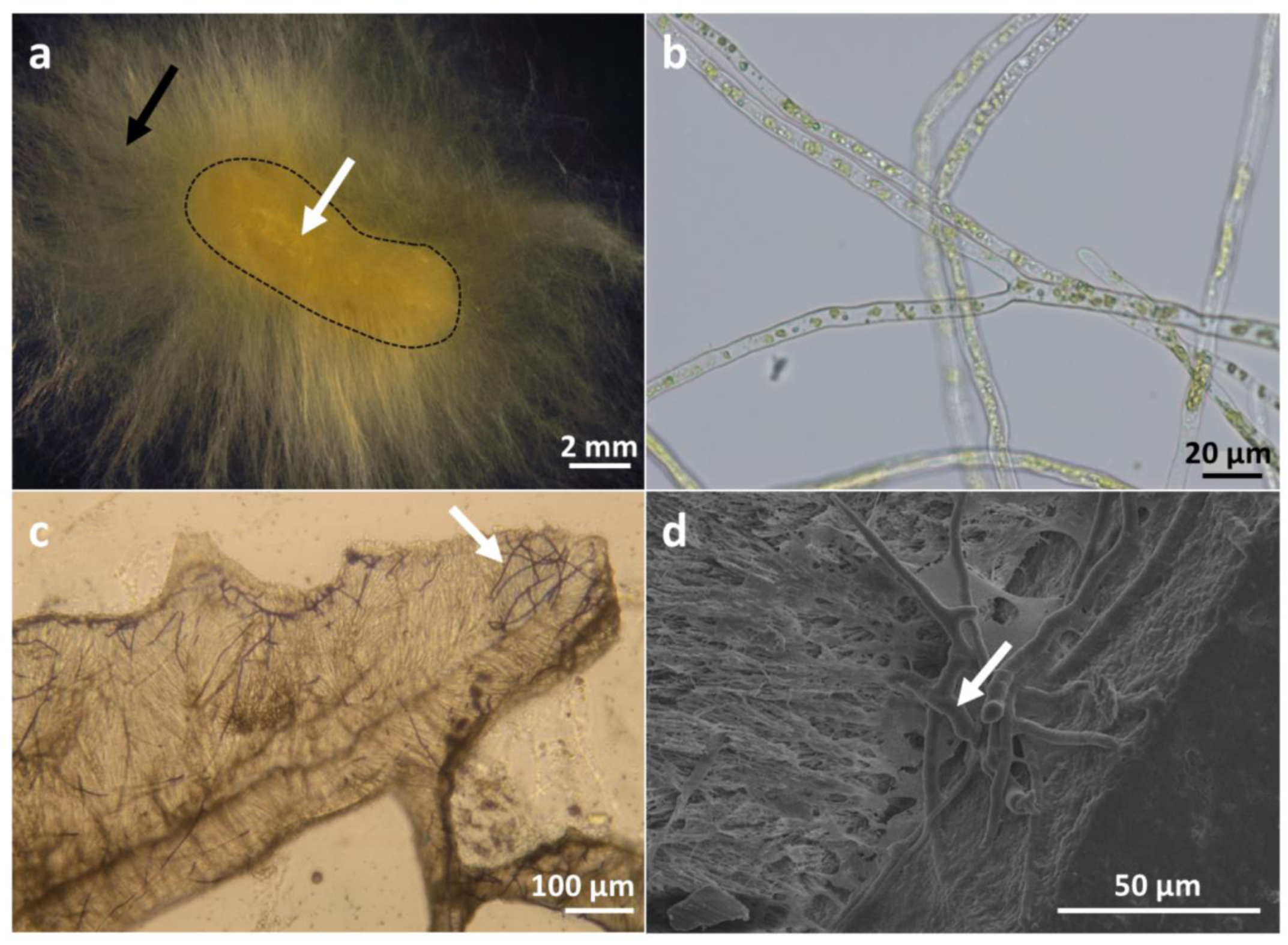
Morphology of *Ostreobium* cultured as free-living or endolithic filaments. Light (a, b, c) and scanning electron (d) microscopy observations of (a) free-living tuft of algal filaments attaching to a fragment of coral carbonate skeleton (outlined by black dotted lines) as epilithic growth form (black arrow) and colonizing this carbonate substrate as endolithic growth form (white arrow). (b) Isolated free-living filaments with visible green chloroplasts inside the branched siphons. (c) Endolithic (bioeroding) filaments (white arrow) stained with toluidine blue in skeleton thin section. (d) Galleries formed by endolithic filaments of *Ostreobium* (white arrow) after re-colonization by free-living filaments of a pre-bleached coral skeleton substrate.

The algal isolates were assigned to 3 species-level *Ostreobium* genotypes (clades) defined by 99% sequence similarity thresholds of the chloroplast-encoded RuBisCo large subunit *rbc*L gene (see Genbank Accession numbers in Table 1). The *rbc*L phylogeny showed that the majority (7/9) of *Ostreobium* strains (obtained from 2 out of 3 host coral colonies) clustered into one P1 clade (99% similarity over ∼729 nt for 6 strains, and over 375 nt for strain 022). Clade P1 is the dominant *Ostreobium* lineage detected in *Pocillopora acuta* corals from long-term cultures at the ATPD-aquarium, and is also detected in *Pocillopora verrucosa* from the Red Sea Eilat IUI reef (Massé *et al*., 2018). The two other strains, named 018A and 06 (each isolated from a single coral colony) were assigned to distinct *rbc*L clades (OTU>99%), named P12 and P14, respectively. Thus, although one coral colony did not provide emerging *Ostreobium* filaments, each of the two other colonies harbored two co-occurring *Ostreobium* lineages, with over-representation of one dominant (P1) genotype.

A phylogenetic tree was built from alignment of overlapping *rbc*L sequences (161 nt length barcode) to assess the strains’ diversity (Fig. S1). Congruent phylogenetic trees and lineage affiliations were also obtained for longer *rbc*L sequence alignments, built over 729 nt for strains 05, 010, 019, 018A, 018B, 018C, and 346 nt for strains 06 and 022 (data not shown). Both clades P12 and P14 were putative sister species-level entities (OTU>99%), related at genus level (>97% sequence similarity threshold) to a clade P3 previously reported in aquarium-grown *Pocillopora acuta* colonies (Massé *et al*., 2018; ATPD and Océanopolis aquariums), and also detected in Pacific reef-collected *Pocillopora* sp. (from Gambier Archipelago, Massé, pers. obs.). Indeed, sequence divergence between P12 and P3, and between P14 and P3 was ∼2%. Both P12 and P14 *rbc*L genotypes clustered together with formerly detected clade P3 into a genus-level lineage (OTU>97%), while P1 *rbc*L genotype was treated separately, as a distinct species-level lineage (OTU>99%). The overall *rbc*L sequence divergence between those lineages was ∼8-10%, with P1 genotype 91.9% similar to P12 genotype and 90.6% similar to P14 genotype.

For comparison purposes with *tuf*A-based *Ostreobium* classification, sequences of the taxonomical marker *tuf*A were also retrieved from available chloroplast genomes of the reference strains *Ostreobium* sp. HV05042 (KY509314) and HV05007a (KY509315) isolated from *Diploastrea* corals (Verbruggen *et al*., 2017), which *rbc*L sequences best matched with the sequences of our strains (Fig. S1). These *tuf*A reference sequences aligned with 97.9% similarity (over 484 nt) to the *tuf*A sequence of *Ostreobium* sp. TS1408 (KU362015) which was affiliated to the family Odoaceae (Sauvage *et al*., 2016), awaiting formal description. This result indicates that the *Ostreobium* strains in P12/P14 lineage were affiliated to the (*tuf*A) Odoaceae family (OTU>92%), while P1 strains represented a species-level genotype belonging to a potentially different family.

### Bioerosive potential in culture conditions

Free-living *Ostreobium* filaments (Fig. 1a, b) were exposed during several weeks to coral skeletal chips measuring a few millimeters in thickness and prepared from pre-bleached *Pocillopora acuta* (hypochlorite cleaned to remove external organic matter). The *Ostreobium* filaments showed attachment to the skeleton surface (epilithic growth habit; Fig. 1a) and entry of some algal filaments perpendicular to the substrate surface (endolithic growth habit). This colonization by endolithic *Ostreobium* was observed by light microscopy on thin sections (Fig. 1c). Toluidine blue-stained filaments penetrating the substrate formed a thick branching network inside the carbonate (Fig. 1c). Galleries formed by this microboring alga (microboring traces called *Ichnoreticulina elegans*; Radtke, 1991) were also observed by scanning electron microscopy (Fig. 1d), below the skeletal surface, confirming *de novo* colonization of the exposed carbonate by endolithic *Ostreobium*. Filaments penetrated through all coral skeletal structures, i.e. growing across bundles of skeletal fibers and centers of calcification without obvious preference for either microstructure. These observations prove that the *Ostreobium* isolates kept their bioerosive potential *in vitro*, even after propagation through several subcultures as free-living forms.

### Photosynthetic and accessory pigments

Qualitatively, the pigment composition of *Ostreobium* strains belonging to lineage P1 (strains of less frequent lineage P12/P14 were not studied) was similar for endolithic and free-living forms (3 and 5 individual strains, respectively, see Table 1). Indeed, similar photosynthetic (chlorophyll) and accessory (carotenoid) pigments were detected for both growth forms in high-performance liquid chromatography (HPLC) fingerprints of organic extracts, although their relative abundance varied (Fig. S2b; with mean retention times (±Standard Deviation) for the 58 peaks detected provided in Table S1). In contrast, profiles of control bleached skeletons (uncolonized by endolithic *Ostreobium*) contained only three peaks, rarely observed and at very low intensity in endolithic *Ostreobium* (Table S1).

Comparison with pure commercial standards allowed identification in endolithic and free-living *Ostreobium* of chlorophyll *a* (chl *a*) in the profiles recorded at 664 nm at retention time (RT) 37.2±0.5 min (with chl *a* allomers at RT 34.6±0.5 min, RT 36.6±0.4 min and RT 38.8±0.5 min) (Fig. S2a). Chlorophyll *b* (chl *b*) was also detected in the profiles recorded at 470 nm, at retention time 30.5±0.5 min (with chl *b* allomers at RT 27.9±0.4 and RT 32.3±0.5 min) (Fig. S2a).

The non-metric multidimensional scaling (nMDS) analysis of profiles showed a great variation of pigment content across strains and subcultures for each *Ostreobium* growth form. Despite this variation, nMDS showed a separation between endolithic and free-living *Ostreobium* phenotypes (except for the free-living subculture 018B^1^ which was close to its corresponding endolithic form) (Fig. 2a2, stress=0.06), with an average dissimilarity of 53% (SIMPER analysis). This dissimilarity between phenotypes was attributed to differences in their photosynthetic pigment proportions, i.e. chl *a*, chl *b*, an allomer of chl *b* (RT 27.9±0.4 min) and two allomers of chl *a* (RT 34.6±0.5 and 36.6±0.4 min), with relative contribution of 14.6%, 13.8%, 10.9%, 7.6% and 5.6%, respectively. Pigment content of both *Ostreobium* growth forms were distinct from the extracts of control bleached skeletons (Fig. 2a1, stress=0.01). A PCA analysis of pigment relative content (not illustrated) showed congruent data point distribution of strains, separating endolithic *versus* free-living phenotypes (except for 018B^1^), with chl *a* and chl *b* and their allomers as main drivers of the distribution pattern.

**Figure 2:**
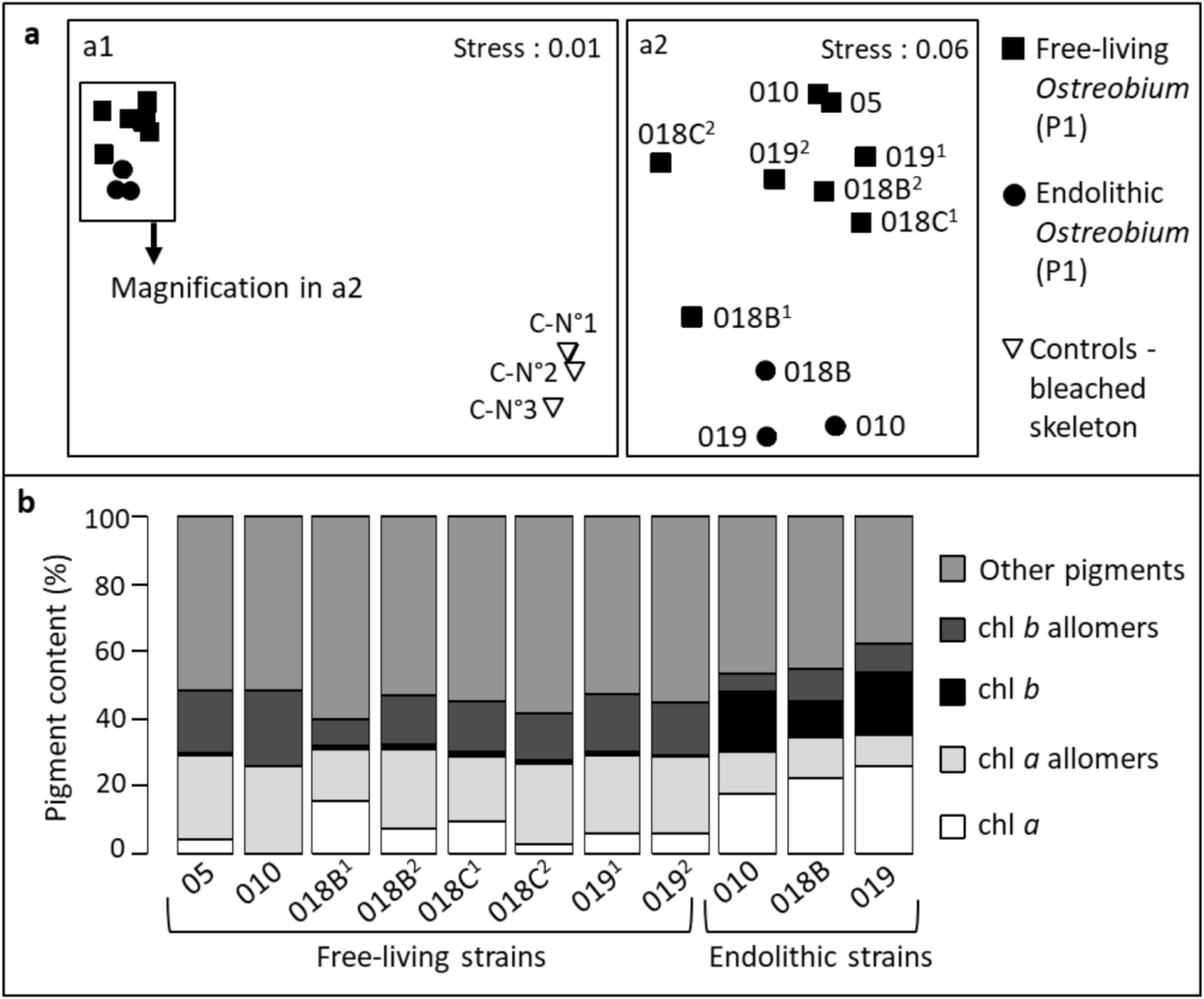
Compared pigment composition of endolithic *versus* free-living *Ostreobium* strains. (2a) Non-metric Multidimensional Scaling (nMDS) plot of Bray-Curtis similarities of pigment composition in endolithic (circles) *versus* free-living (squares) *Ostreobium* strains (named 0xx) belonging to lineage P1 (in black), and control bleached coral carbonate skeleton (triangle in white). ^1^ and ^2^ indicate successive subcultures of strains. (2b) Barplot of relative proportions of pigments in individual strains (P1) and growth forms.

Quantitatively, a high inter-individual variability was noted for chlorophyll content (chl *a* and chl *b*, and their respective allomers) especially across free-living strains and their successive subcultures (illustrated in Fig. 2b and detailed in Table S2). Overall, despite variable chl *a* content between subcultures of the free-living phenotype, chl *b* contents were higher in endoliths (8.9±1.2 µg/mg organic extract, respectively; SE, n=3) than free-living forms (1.0±0.3 µg/mg organic extract, respectively; SE, n=8). The ratio chl *b* : chl *a* was four times higher in endolithic (0.84±0.18, SE, n=3) than in free-living (0.21±0.03, SE, n=8) *Ostreobium* (Table S2).

### Fatty acid composition

Qualitatively, the fatty acid (FA) composition of *Ostreobium* strains belonging to P1 and P12/P14 lineages was similar between endolithic and free-living forms (Table S3). Representative fatty acid profiles obtained by gas chromatography (GC) are illustrated in Figure S3. A total of 31 FAs were detected in *Ostreobium* strains, dominated by saturated fatty acids (SFA) in endoliths *versus* polyunsaturated fatty acids (PUFA) in free-living forms (Table S3). In contrast, only 5 saturated FAs (16:00, 18:00, 14:00, 15:0 and 12:0) were found in control bleached skeletons (detailed in Table S3).

Quantitatively however, PCA and barplot analysis showed variability of FA content (relative proportions) across *Ostreobium* strains and subcultures for each phenotype (Fig. 3). Despite this variability, a general trend could be observed and showed a different FA content in *Ostreobium* strains belonging to clade P1 (black color code, Fig. 3a) than in the majority of their free-living counterparts (for all cultures but 010, 018B^1^, 019, see also Fig. 3c). For clade P12/P14, due to low number of available strains and limited available endolithic biomass, the recorded similarity of FA content of endoliths and free-living forms is provisional because it was based on a single determination for endolithic strain 06 (see Table S3).

**Figure 3:**
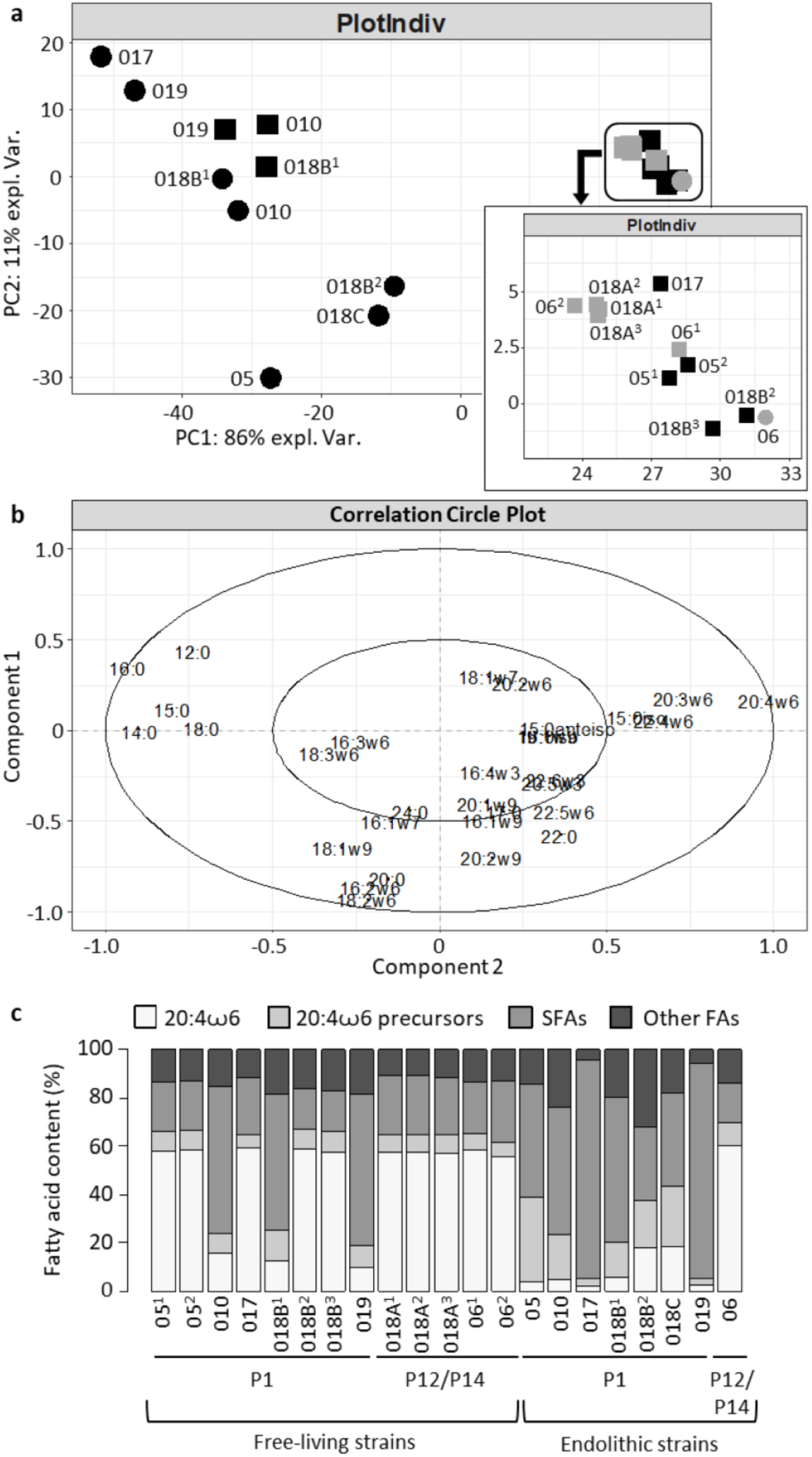
Fatty acid composition of endolithic *versus* free-living *Ostreobium* growth forms. (a) Scores plot and (b) loading plot for PCA analysis of relative fatty acid methyl ester (FAME) proportions measured in GC profiles after GC-MS annotation of endolithic (circle) and free-living (square) strains (named 0xx). Genetic lineages are color coded (black: P1; grey: P12/P14). (c) Barplot of relative proportions of fatty acids in individual strains and growth forms. Means and Standard Error (SE) of % total fatty acids are detailed in Table S3, along with polyunsaturated and saturated fatty acid values per growth form and lineage.

The poly-unsaturated fatty acid content explained 86% of the variability between endolithic and free-living phenotypes (PC1, Fig. 3a, 3b). Indeed a major difference was due to arachidonic acid (20:4ω6) content increased by a factor ranging from 2.2 to 30 in P1 lineage, from endolithic to free-living forms (illustrated in Fig. 3c, see also Table S3; a large range attributed to variable physiological status across subcultures). Biosynthetic intermediates of arachidonic acid, i.e. 16:3ω6, 18:2ω6 and 18:3ω6 were detected in endolithic *Ostreobium*, like in free-living forms, but without longer-chain 20:4ω6 fatty acid. Among detected mono-unsaturated fatty acids, 18:1ω7 was abundant in both *Ostreobium* forms, as well as 18:1ω9 which was simultaneously detected with 18:2ω6. Trace amounts (<0.5%) of methyl-branched fatty acids (15:0iso, 15:0anteiso and 16:0iso) and long-chain PUFAs (20:5ω3, 22:5ω6 and 22:6ω3) were also detected in both *Ostreobium* forms, in variable amounts across strains and their subcultures.

### C/N ratios and stable isotope values (δ^13^C and δ^15^N)

The carbon to nitrogen (C/N) ratios in endolithic *Ostreobium* filaments removed from their surrounding carbonate (by acid-decalcification), did not vary much among strains and subcultures and between lineages, with mean C/N of 12.6±0.6 for lineage P1 (n=5, SD) and 11.9±0.3 for lineage P12/P14 (n=4, SD) (Table S4). These values are close to those measured in the residual organic matrix from control acid-decalcified bleached skeletons (mean C/N ratios of 11.1±1.1, n=4, SD). Conversely, in free-living forms, the C/N ratio varied greatly among strains within both P1 and P12/P14 lineages (C/N of 16.3±2.5 n=9, SD for P1, and 14.3±1.4 n=4, SD for P12/P14), with more stable ratios between subcultures of each strain. The overall pattern indicates that C/N ratios were higher in free-living than in endolithic filaments of *Ostreobium*.

Stable isotope values (δ^13^C and δ^15^N) of endolithic (not decalcified, or acid-decalcified) and free-living *Ostreobium* forms are presented in Figure 4 along with controls, i.e. bleached coral skeleton substrate and its residual organic matrix. Detailed δ^13^C and δ^15^N values of individual strains and growth forms are also presented in Table S5. Results showed high δ^13^C variability among strains and their subcultures, within and between each genetic lineage (P1 and P12/14), for each growth form. Despite this variability, data of decalcified endolithic *Ostreobium* were pooled as mean δ^13^C values were not significantly different between lineages (p>0.05). For the same reason, data of free-living *Ostreobium* were also pooled (Fig. 4A). The non-decalcified endolithic *Ostreobium* belonging to P1 (non-decalcified P12/P14 were not measured) were significantly (p<0.05) more enriched in ^13^C (−11.2±0.8 ‰ SD, n=4) than the free-living phenotypes (−18.6±2.6 ‰ SD, n=16 for all clades). In contrast, decalcified endolithic *Ostreobium* were significantly (p<0.05) more depleted in ^13^C (δ^13^C −23.8±3.6 ‰ SD, n=13 for all clades) than the free-living forms. This depletion is partly explained by the acid treatment of endolithic filaments to remove the carbonate. Indeed a parallel control test of this acid treatment on free-living *Ostreobium* filaments showed a significant (p<0.05) −6 ‰ amplitude depletion of δ^13^C (i.e. −26.9±2.6 ‰ SD, n=3 treated *versus* −20.9±2.6 ‰ SD, n=3 untreated, technical replicates of strain 010, Table S5). Thus, acid treatment corrected δ^13^C values of decalcified endolithic *Ostreobium* strains were on average −17.8±3.6 ‰ (SD, n=13 for all clades). However, when strain 010 was excluded from the pool of data as it seems to behave differently from the other strains, decalcified endolithic strains still appeared depleted in ^13^C (corrected δ^13^C : −20.1±2.3 ‰ SD, n=8, for all strains except 010) compared to their corresponding free-living form (see Table S5).

**Figure 4:**
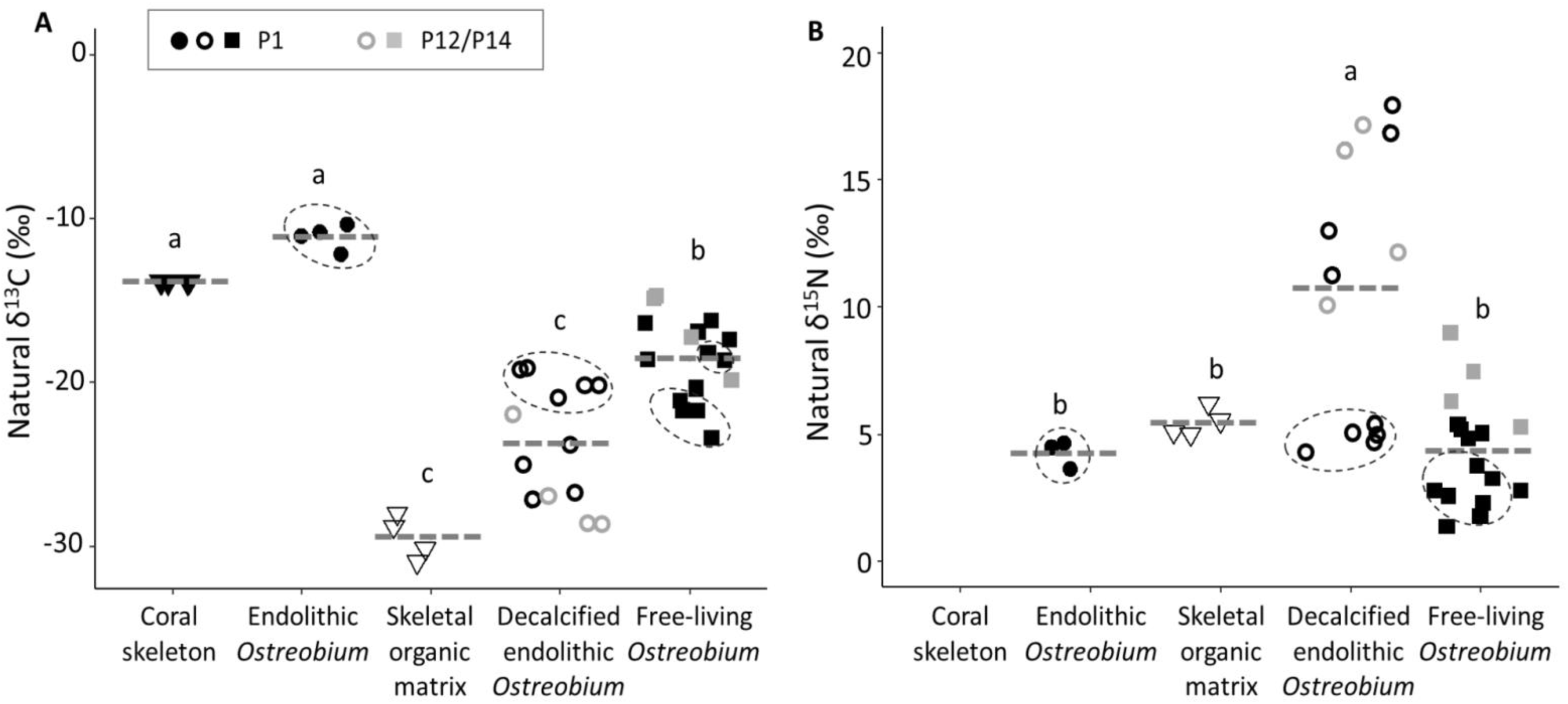
Stable isotope values (δ^13^C and δ^15^N) of endolithic *versus* free-living *Ostreobium*. Endolithic filaments were analyzed by EA-IRMS either within their carbonate substrate (endolithic *Ostreobium*, filled circles) or after decalcification with formic acid 5% (decalcified endolithic *Ostreobium*, empty circles). The acid treatment depletes δ^13^C values of decalcified endolithic *Ostreobium* by −6 ‰ (here data are presented before correction). Control substrate of endoliths was either bleached coral carbonate skeleton (composed of CaCO_3_ and organic matrix, filled triangle) or skeletal (acid-insoluble) organic matrix (empty triangles). Corresponding free-living filaments (squares) were also analyzed. Genetic lineages are color coded (black: P1; grey: P12/P14). Intra-carbonate δ^15^N of coral skeleton was below EA-IRMS detection limit. For graph clarity, strain code names are not indicated, except for dotted circled δ^13^C and δ^15^N values of strain 010 which plot separately from other P1 strains. Grey bars indicate means. Different letter indicate significant differences (ANOVA and student test p<0.05). δ^13^C and δ^15^N values of individual strains and growth forms are provided in Table S5.

The mean δ^13^C of endolithic *Ostreobium* was compared to that of their habitat (coral skeleton substrate) to track shifts indicative of potential C sources. Results showed that the δ^13^C values in undecalcified endolithic *Ostreobium* were higher (−11.2±0.8 ‰ SD, n=4 clade P1 strain 010) than those of control skeleton substrates (−13.9±0.03 ‰ SD, n=3). Such ^13^C enrichment (+2.7 ‰) was also observed and was significant (p<0.05) using another sample preparation method, the automated carbonate preparation device Kiel IV (endolithic *Ostreobium* δ^13^C −10.6±0.4 ‰ SD, n=8 clade P1 strain 010 *versus* carbonate substrate δ^13^C −13.3±0.4 ‰ SD, n=8). Regarding decalcified endolithic *Ostreobium*, their δ^13^C values were higher (−23.8±3.6 ‰ SD, n=13) than those of their control decalcified skeleton substrate, i.e. the coral skeletal organic matrix (−29.4±1.3 ‰ SD, n=4).

For δ^15^N (Fig. 4B), similarly to δ^13^C, high inter-individual variability was observed for each growth form, between strains and their subcultures and within each genetic lineage (P1 *versus* P12/14). However, this variability was much higher for the decalcified endoliths (+4.3 ‰ min, +17.9 ‰ max), than for the free-living forms (+1.4 ‰ min, +9.0 ‰ max). The lineage had a significant effect on mean δ^15^N of free-living *Ostreobium* (p>0.05) but not on that of decalcified endoliths (due to high variability within lineage). In order to compare N sources between growth forms, data for both genetic lineages were pooled. Results showed that the δ^15^N values of most decalcified endolithic *Ostreobium* strains were significantly enriched (14.3±3.1 ‰ SD, n=8 for all clades, except strain 010) compared to the δ^15^N values of their corresponding free-living forms (5.3±2.1 ‰ SD, n=10 for all clades, except strain 010). Only for strain 010 within P1 (dotted circles in Fig. 4B), the recorded δ^15^N values were low and similar for both the non-decalcified and decalcified endolithic form (4.2±0.6‰ SD, n=3 and 4.9±0.4‰, SD, n=5, respectively), but still enriched compared to the δ^15^N of their free-living counterparts (2.7±0.7‰ SD, n=6). There was no effect of the acid treatment as δ^15^N values of control treated or untreated free-living forms were not significantly different (2.2±1.2 ‰ SD, n=3 treated *versus* 2.3±0.5 ‰ SD, n=3 untreated, technical replicates of strain 010; Table S5). Compared to δ^15^N of the coral skeletal organic matrix (5.5±0.5 ‰ SD, n=4), most values of the decalcified endolithic *Ostreobium* strains (except strain 010) were enriched (14.3±3.1 ‰ SD, n=8) corresponding to a +9‰ average shift.

### Photosynthetic C and N assimilation

Bicarbonate and nitrate uptake by endolithic *Ostreobium* and their corresponding free-living forms (n=3 strains of P1 lineage, and n=2 strains of P12/P14 lineage) are illustrated in Figure 5, in relation to pH changes during the 8h dual labeling pulse with ^13^C-bicarbonate and ^15^N-nitrate, under light or dark conditions. Different patterns of C and N assimilation were recorded between endolithic *versus* free-living phenotypes, despite strain-related variations. Indeed a high metabolic variability was observed among strains within each lineage (P1 or P12/P14). Due to low replication, the lineage effect could not be tested (experimental culturing difficulties limited to 2 the number of available P12/P14 strains and strain 010 in P1 was analyzed only in light conditions).

**Figure 5:**
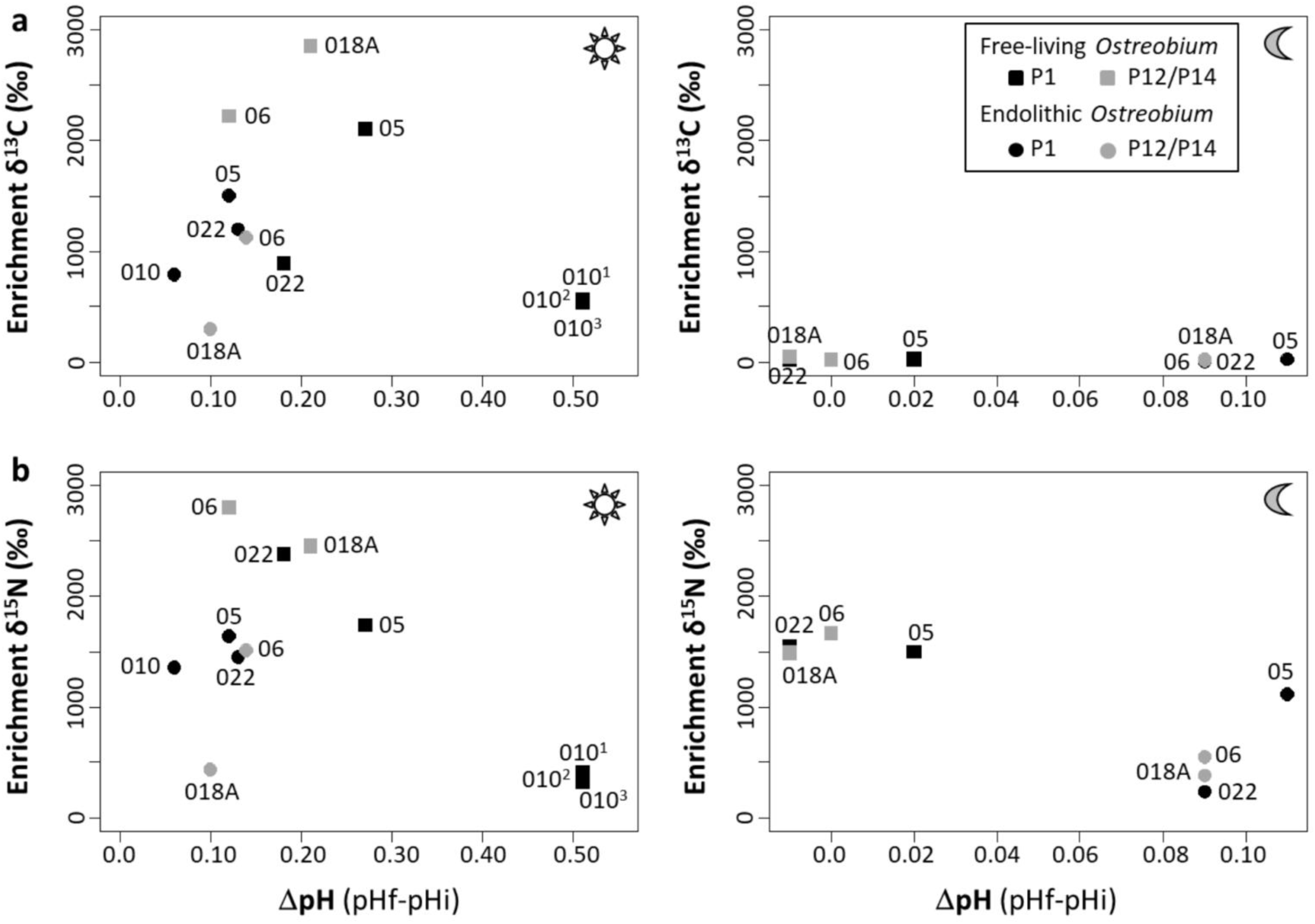
Variability of photosynthesis-dependent inorganic carbon and nitrogen assimilation among endolithic *versus* free-living *Ostreobium* strains in relation to pH changes. Values of (a) enriched δ^13^C and (b) enriched δ^15^N in free-living and decalcified endolithic strains were calculated at the end of the 8h labeling pulse with ^13^C-bicarbonate (2 mM) and ^15^N-nitrate (5 µM) in light or dark conditions, via comparison to corresponding unlabeled controls. ΔpH=pHf-pHi, with final pHf at the end of 8h labeling pulse compared to initial pHi. Genetic lineages are color coded (black: P1; grey: P12/P14), and strain code names (0xx) are indicated. ^1^ and ^2^ indicate technical replicates of strains 010.

Fixation of ^13^C-bicarbonate (Fig. 5a) in light condition was lower in endolithic forms (973±616 ‰ SD, n=3) than in free-living forms (2392±405 ‰ SD, n=3) for three strains: 05 (P1) and 06, 018A (P12/P14). However, strains 022 and 010 (both in P1) displayed an opposite pattern. In dark conditions, very little ^13^C enrichment was recorded for both endolithic (14.9±10.2 ‰ SD, n=4) and free-living *Ostreobium* (30.5±11.6 ‰ SD, n=4) for all strains, indicating that ^13^C enrichment resulted from photosynthesis. Indeed, in light conditions photosynthetic activity was confirmed by the rise in pH during the 8h labeling pulse, for both endolithic (+0.11±0.03 SD, n=5) and free-living *Ostreobium* (+0.26±0.15 SD, n=5). For endolithic phenotypes, seawater alkalization may also result from carbonate dissolution activity, in addition to photosynthesis. Although the pH increase was much lower in endolithic than free-living forms (consistent with lower biomass of filaments: 0.9±0.4 mg *vs* 5.9±2.6 mg dry weight, n= 5, respectively), large pH increase corresponded to high ^13^C-enrichment, except for the free-living form of strain 010 (clade P1). In dark conditions, pH did not vary for the free-living forms while it increased for the endolithic forms (+0.1±0.01 SD, n=4), indicative of carbonate dissolution processes.

Assimilation of ^15^N-nitrate (Fig. 5b) was recorded at the end of the 8h labeling pulse in both endolithic and free-living *Ostreobium* in light and also in dark conditions. Uptake of ^15^N-nitrate was however 40% reduced in dark compared to light conditions. In light, assimilation of ^15^N-nitrate was lower in endolithic forms (1253±556 ‰ SD, n=4) than in free-living forms (2346±442 ‰ SD, n=4) for four strains (06, 018A, 022, 05). Strain 010 displayed an opposite pattern. In dark conditions, ^15^N-assimilation was also lower in endolithic forms (567±384 ‰ SD, n=4) than in free-living forms (1552±81 ‰ SD, n=4).

We calculated the turnover of exogenous inorganic ^13^C and ^15^N (i.e. the rate of inorganic C and N used for cell growth) in endolithic *versus* free-living filaments for a 12h photoperiod, by extrapolation of the uptake data recorded at the end of the 8h pulse of ^13^C-bicarbonate and ^15^N-nitrate (equation in Material & Methods and data for each strain in Table S6). In daylight, C and N turnover rates varied among strains for both phenotypes. Although variability between technical replicates within a same strain was low (SD<5% of the mean for free-living strain 010), differences were observed between genetic lineages (Table S6). Indeed, C and N turnover increased by a factor ∼3 in P12/P14 compared to P1 when *Ostreobium* filaments lived free in the culture medium, whereas P12/P14 was less active than P1 when filaments lived inside carbonate substrate (with C and N turnover ∼60 and 50 % weaker, respectively). Regarding phenotypes, C and N turnover in P1 were relatively similar between growth forms, whereas in P12/P14 it decreased by a factor 3.6 for C, and a factor 2.7 for N in endoliths compared to free-living *Ostreobium* forms. In dark conditions, C turnover for both phenotypes was very low (<0.1 % / 12h period, Table S6). In contrast, N turnover was detected for both phenotypes but at reduced level compared to light conditions. For free-living *Ostreobium*, N turnover was similar among strains and between genetic lineages (Table S6). For endolithic *Ostreobium*, N turnover greatly varied among strains within lineage P1, whereas lower variability was observed within P12/P14 (Table S6). Regarding phenotypes, N turnover decreased in endolithic compared to free-living *Ostreobium*, by a factor 2.2 for lineage P1 and 3.3 for lineage P12/P14. Combined together, these C and N turnover results indicated a lower efficiency of dissolved inorganic carbon and nitrogen (DIC & DIN) uptake from seawater by endolithic *Ostreobium* than by their free-living counterparts.

## Discussion

### Diversity of *Ostreobium* strains from *Pocillopora* corals

Despite rapidly growing evidence of the high genetic diversity of the microboring Chlorophyta *Ostreobium* in living corals in reef ecosystems (Marcelino & Verbruggen, 2016; Sauvage *et al*., 2016; Marcelino *et al*., 2017; del Campo *et al*., 2017; Gonzalez-Zapata *et al*., 2018), phenotypic characterization of specific *Ostreobium* genotypes has lagged behind. Indeed, phenotype studies have so far focused only on strains 6.99 and B14.86 designated as *Ostreobium quekettii* Bornet & Flahaut 1889 (Kornmann & Sahling, 1980; Schlichter *et al*., 1997; Krause *et al*., 2019). Phenotypic comparisons of *Ostreobium* strains belonging to distinct genetic lineages are key to better understand the functional diversity of this microboring alga in reef ecosystems. Here, the nine *Ostreobium* strains obtained from the fast-growing branching coral *Pocillopora acuta* Lamarck 1816 add to the diversity of the few coral-isolated strains (mostly from massive, slow-growing coral hosts) that were so far used only for phylogenetic studies (Verbruggen *et al*., 2017; Sauvage *et al*., 2016). Although isolated from aquarium-grown, long-term propagated coral colonies, these *Ostreobium* strains are representative of 2 lineages from reef settings, including a lineage provisionally assigned to the family Odoaceae (*sensu* Sauvage *et al*., 2016), which are also detected in natural communities of *Pocillopora* corals from Northern Red Sea and South Pacific reefs (Massé *et al*., 2018; Verbruggen *et al*., 2017).

Fast-growing branching corals are known to harbor less abundant and diverse *Ostreobium* than slow-growing massive corals, with patchy spatial distribution along the branch growth axis (Godinot *et al*., 2012; Massé *et al*., 2018; Marcelino *et al*., 2017). Culture bias likely allowed only a fraction of endolithic microbes to be isolated and potential multiple isolation of the same *Ostreobium* lineage due to filament fragmentation, with one dominant (P1) and one minor (P12/P14) lineage detected per branch. These reasons explain the uneven size of datasets generated in this study, largest for strains affiliated to the P1 lineage (7 strains), and reduced to 2 strains for the P12/P14 lineage. Isolation and characterization of more strains from P12/P14 would be necessary to further test metabolic differences between lineages. Nevertheless, this collection of *Ostreobium* strains that kept their bioerosive potential after isolation from a branching *Pocillopora* coral species, provides novel tools to study the nature of *Ostreobium* interactions with branching corals and their endosymbiotic *Symbiodiniaceae*, sensitive to bleaching events in a changing environment.

### Habitat-driven changes in morpho-functional traits (pigments and fatty acids)

Pigments of *Ostreobium* algae have so far been mainly studied in complex natural microboring communities, i.e. green bands of dead or live corals, using spectrophotometry and Jeffrey and Humphrey equations for chlorophylls, or HPLC technique (Fork & Larkum, 1989; Fine & Loya 2002; Fine *et al*., 2006; Tribollet *et al*., 2006; Sangsawang *et al*., 2017) with a few pioneer studies focusing on individual strains of *Ostreobium* sp. (called *O. quekettii* Bornet & Flahaut 1889) using HPLC methods (Jeffrey, 1968; Schlichter *et al*., 1997; Koehne *et al*., 1999). For the first time, we provide pigment datasets (HPLC profiles) for several *Ostreobium* strains of a referenced genotype (*rbc*L P1 lineage) and show differences in endolithic *versus* free-living forms, supporting adaptation to habitat-driven lower light intensities in the dense coral biomineral *versus* seawater. Indeed, although pigment composition was qualitatively similar among strains, chlorophyll *b* content and chl *b* : chl *a* ratio were higher in endolithic forms compared to free-living forms. These results obtained for genetically identified *Ostreobium* strains are in agreement with those reported for natural microboring communities, highlighting the major contribution of *Ostreobium* to the photobiology of complex coral skeletal microbiomes. Indeed, in the green layer of massive (*Favia*) corals, chlorophyll *b* reached up to 60-75% of the chlorophyll *a* content (Jeffrey, 1968). In the endolithic algae colonizing the coral *Leptoseris frugilix*, the chl *b* : chl *a* ratio was also higher in deep colonies than shallow ones (Schlichter *et al*., 1997). Additionally, in free-living cultures of *Ostreobium* sp. strain B14.86, Schlichter *et al*. (1997) experimentally showed that chlorophyll *a* and *b* concentrations increased with decreasing light (photon flux from 60 to 0.5 µmol.m^-2^.s^-1^), suggesting adaptation of the light harvesting system to low light conditions. Our experimental results indicate that endolithic *Ostreobium* strains adapted their photosynthetic apparatus (chl *b* content) to the low light microenvironment inside the dense coral carbonate substrate (Enriquez *et al*., 2017), which here was the bleached skeleton (bared of tissue) of a fast-growing, branched coral. Contrary to other studies on (unidentified) *Ostreobium* sp. strain from Chilean coral (Koehne *et al*., 1999; Wilhelm & Jakob, 2006), or complex endolithic community from coralline algae (Behrendt *et al*., 2011), the far-red absorbing chlorophyll *d* was not detected (i.e. no peak with a maxima absorption around 700 nm) in our endolithic *Ostreobium* strains. Red-shifted chlorophyll *d* is mainly associated to photoadaptation of *Ostreobium* algae to life in low light microenvironment inside the coral carbonate skeleton of living colonies, shaded by intact coral tissue cover (Haldall, 1968). Our experimental settings, i.e. illumination with ∼30 µmol photons.m^-2^.s^-1^ white light and absence of light-absorbing coral tissue cover, may have prevented the full expression of *Ostreobium* pigment repertoire. Future studies may involve manipulation of light quality and intensity, and use of high spatial resolution sensors (Magnusson *et al*., 2007; Wangpraseurt *et al*., 2012), to better resolve the optical microniche environments inside coral skeletons *versus* inside free-living tufts of filaments, and compare the photobiology and photoadaptation of diverse genotypes of *Ostreobium*.

Fatty acids are important for physiological adaptation of algae to environmental stress such as salinity and temperature (Zhila *et al*., 2010, de Carvalho & Caramujo, 2018). They may also be used to understand the nutritional value of *Ostreobium* to the coral host (suggested by Fine & Loya, 2002) and to grazing organisms such as parrotfishes (Clements *et al*., 2016), and more generally, as trophic markers to trace food sources in reef organisms. Here we provide first detailed records of *Ostreobium* fatty acid profiles, highlighting their variability across growth forms, genotypes, strains and subcultures.

Fatty acid composition of *Ostreobium* was typical of chlorophytes but varied quantitatively between the endolithic growth forms and their free-living counterparts. Palmitic acid (16:0), abundant in both endolithic and free-living *Ostreobium*, is indeed a saturated FA reported in several green macroalgae (Kumari *et al*., 2010; Pereira *et al*., 2012). Both phenotypes displayed high contents of C18 polyunsaturated FA (PUFA) which are characteristic of chlorophytes (Jamieson & Reid, 1972; Khotimchenko & Svetashev, 1987). But some FAs commonly found in other algae of the order Bryopsidales were not detected in our *Ostreobium* strains. Indeed, a comparison of Ulvales and Cladophorales with the order Bryopsidales (the latter listed by Aknin *et al*. (1992) as Siphonales, a previous affiliation of members of this order) indicates that these algae are characterized by high quantity of 16:3ω3 combined with low quantities of 16:4ω3 and 18:4ω3 (Aknin *et al*., 1992). The absence of these ω3 fatty acids from all analyzed *Ostreobium* strains suggests their absence in the suborder Ostreobinae, at least in the two lineages to which our strains were classified.

Simultaneous detection in our *Ostreobium* cultures of 18:2ω6 and 18:1ω9 is indicative of fungal presence (Frostegård *et al*., 1996; Mikola & Setälä, 1998; Chen *et al*., 2001; Meziane *et al*., 2006), and confirms that cultures were not axenic. Fungi from the microbiome of the coral skeleton may have been co-isolated. Structures morphologically similar to fungal hyphae are indeed known to colonize *Ostreobium* filaments in various carbonate substrates (Le Campion-Alsumard *et al*., 1995b; Tribollet & Payri, 2001; Golubic *et al*., 2005). Likewise, trace amounts of methyl-branched saturated fatty acids were detected, indicative of bacterial presence (Daalsgard *et al*., 2003). Endophytic bacteria have indeed been reported in the genus *Bryopsis* (Hollants *et al*., 2011) and some bacteria may be associated with *Ostreobium* filaments. Long-chain PUFAs (20:5ω3, 22:5ω6 and 22:6ω3) typical of dinoflagellates (Mansour *et al*., 1999) were also sometimes detected at low levels, especially in some of the free-living forms.

An important feature of the fatty acid composition of *Ostreobium* strains was the high abundance of arachidonic acid (20:4ω6) in free-living forms, consistently dropping in endolithic forms (observation valid for lineage P1, as low replication of endolithic forms in the less represented P12/P14 prevented comparisons). This essential FA for living organisms is a constituent of phospholipids in biological membranes, involved in fluidity, permeability and cellular signalization (Maulucci *et al*., 2016). The shift recorded in *Ostreobium* membrane composition suggests important adjustments of fluidity and permeability, possibly for adaptation to the lifestyle constraint experienced by the siphons. Endolithic *Ostreobium* siphons are indeed supported by the rigid environment provided by the coral carbonate mineral. C/N ratios were also lower in endolithic *versus* free-living *Ostreobium* suggesting lower production inside the carbonate habitat of C-rich compounds such as lipids and polysaccharides exudates. In contrast, free-living filaments may benefit from more fluid and flexible membranes that offer more mechanical resistance to shear forces and local seawater turbulence. In addition high PUFA content may provide more protection from temperature changes (de Carvalho & Caramujo, 2018). Decreased degree of fatty acid unsaturation in endoliths could also help the alga to acclimate to salinity stress (Zhila *et al*., 2010). Overall a shifting arachidonic acid content between both phenotypes may reflect differential signalization activity (de Carvalho & Caramujo, 2018), and a shift in communication with associated microbes. Such potential differences could occur within the same filament, with one end as free-living *Ostreobium* emerging as epilith from carbonate substrates and the other end as endolith. This remains to be investigated in light of the important nutritional role of epilithic and endolithic biofilms in reef trophic food chain (Kobluk & Risk, 1977; Adey, 1998; Clements *et al*., 2016).

### Nutritional (C and N) sources for *Ostreobium* phenotypes

The sources of carbon (C) and nitrogen (N) used by endolithic *Ostreobium* for photosynthesis and growth inside a coral carbonate habitat remain little known. A few authors showed that microboring communities dominated by *Ostreobium* sp. in dead reef carbonate substrates are stimulated by elevated seawater *p*CO_2_, suggesting that those microboring algae are limited in DIC and use mainly seawater DIC (Tribollet *et al*., 2009; Reyes-Nivia *et al*., 2013; Tribollet *et al*., 2019). A similar trend was also observed for another reef microboring community dominated by the chlorophyte *Phaeophila* sp. in dead carbonate substrates (Enochs *et al*., 2016). In light of a recent study by Guida *et al*. (2017), endolithic microalgae such as *Ostreobium*, could also use bicarbonate ions (HCO_3_^-^) released during carbonate dissolution for photosynthesis. Indeed, these authors showed that the microboring cyanobacterium, *Mastigocoleus testarum* (in cultured strain BC008 and under natural conditions), is able to fix carbon derived from mineral calcite substrate when seawater DIC is limiting. Here, the stable isotope analysis of *Ostreobium* strains provides new information on their sources of carbon and nitrogen for photosynthesis and nutrition.

Regarding carbon, the only possible source of carbon for free-living *Ostreobium* was DIC from seawater, which is confirmed by the uptake of ^13^C-bicarbonate in our experiment (Fig. 5a). Their measured δ^13^C values (ranging between −24 and −15 ‰) were in agreement with those recorded for green fleshy macroalgae (Raven *et al*., 2002), especially in the class Ulvophyceae (Wang & Yeh, 2003). For endolithic *Ostreobium* boring through the coral biomineral, multiple C sources are possible: (i) DIC from seawater and diffusing to the interstitial space at the interface between filament and skeleton; (ii) DIC released from the CaCO_3_ biomineral by active carbonate dissolution and (iii) organic C released from the skeletal organic matrix by carbonate dissolution and/or the activity of microbial associates. Here, we show that non-decalcified endolithic *Ostreobium* filaments (δ^13^C −11.2±0.8 ‰) were depleted in ^13^C compared to reference seawater DIC (in the range of −5 to 2 ‰; see Patterson & Walter, 1994 for δ^13^C values of tropical seawater in carbonate reefs), and more enriched compared to their coral carbonate substrates. Indeed, the mean δ^13^C value of bleached *Pocillopora acuta* coral skeletons was −13.9±0.03 ‰, which corresponds to the low range of values recorded for hermatypic, symbiotic coral skeletons (Linsley *et al*., 2019), likely due to aquarium coral growth restrictions and maybe reduced photosynthesis, increased respiration and oxidation of organic matter. This result, combined with the ^13^C-bicarbonate uptake experiment, strongly supports the hypothesis that endolithic *Ostreobium* used mainly DIC from seawater.

After correction for acidification treatment (+6 ‰), most decalcified endolithic *Ostreobium* strains had lower δ^13^C compositions compared to that of their corresponding free-living forms (Table S5). The uptake of ^13^C-labeled bicarbonate and exogenous C turnover were however reduced (Fig. 5a and Table S6, respectively) in endolithic compared to free-living *Ostreobium*, indicating lower photosynthetic activity inside the light-limited carbonate. This supports the hypothesis that most endolithic *Ostreobium* had access to the same source of seawater carbon as their free-living counterparts. Indeed, low photosynthetic rates are expected to reduce δ^13^C values of macroalgae (O’Leary, 1988; Wiencke & Fischer, 1990) due to preferential uptake of dissolved CO_2_ if not limiting, and carbon isotope fractionation during photosynthetic ^12^C-CO_2_ fixation by the RuBisCO enzyme, which discriminates against ^13^C-CO_2_ (Farquhar *et al*., 1989). Moreover, the presence of contaminant residual skeletal organic matrix with very low δ^13^C values (−29.4±1.3 ‰) also likely contributed to depletion of ^13^C in decalcified endolithic *Ostreobium*.

Only endolithic filaments of strain 010 had enriched δ^13^C values compared to their free-living filaments (Fig. 4, Table S5). This could be explained by the relatively low δ^13^C values of its free-living phenotype due to low photosynthetic activity, supported by very low chlorophyll content (Table S2) and reduced exogenous C turn-over (Table S6). It might also point to an uptake by the endolithic phenotype of strain 010 of non-isotopically labeled DIC, released by dissolution of the coral CaCO_3_ substrate, and carbonic anhydrase conversion of bicarbonate ions (Shashar & Stambler, 1992) which would increase its δ^13^C values. Indeed in dark conditions, we recorded a significant pH increase in endolithic cultures (Fig. 5) suggesting *Ostreobium*-driven carbonate dissolution. Moreover, a preliminary experiment with *Ostreobium* strain 018B also indicated *in vitro* production of alkalinity over a 24h day/night cycle (measured by colorimetry according to Sarazin *et al*., 1999). Combined together, these results suggest that the endolithic *Ostreobium* strains were dissolving the coral carbonate during our experiments. This dissolution is consistent with recent studies that showed for *Ostreobium quekettii* strain (Krause *et al*., 2019) or microboring communities dominated by *Ostreobium* in dead coral skeletons (Tribollet *et al*., 2019) that endolithic filaments are actively dissolving carbonates, increasing seawater alkalinity and thus, the concentration of bicarbonate ions in their environmental vicinity. Although in our experimental settings most endolithic strains were using seawater DIC, the relative contribution of each carbon source for endolithic *Ostreobium* growth forms needs to be further investigated, testing a range of seawater DIC. The possible use by endolithic *Ostreobium* of organic C released from the skeletal organic matrix and the activity of associated microbes should also be investigated. This would allow to better understand *Ostreobium* nutritional plasticity and their efficiency at dissolving carbonates under different environmental conditions.

Regarding nitrogen, the positive shift of δ^15^N values recorded in this study for endolithic compared to free-living *Ostreobium* suggests a higher trophic level and different nitrogen sources. The increased δ^15^N values may also reflect nutrient limitation and complete utilization by endoliths of the nitrate source pool in the small interstitial spaces between *Ostreobium* filaments and their carbonate substrate (Torres *et al*., 2012). For free-living forms, the only N source was dissolved inorganic nitrogen (DIN) in the form of nitrate (NO_3_^-^) from seawater medium, confirmed by uptake of ^15^N-nitrate in our experiment (Fig. 5b). For endolithic forms, two N sources are possible: (i) DIN from seawater and diffusing to the interstitial space at the interface between filament and skeleton and (ii) organic N released from the skeletal organic matrix by carbonate dissolution and/or from the activity of microbial associates. Similarly to the free-living forms, nitrate provided in the incubation seawater was assimilated by *Ostreobium* endoliths, as shown by their ^15^N-enrichment at the end of the 8h pulse of labeling with ^15^N-nitrate. The nitrate uptake was however higher in light than dark conditions. This reduction (40%) is likely explained by the contribution of photosynthesis to supply energy and organic carbon skeletons (C-backbones) for nitrogen assimilation into amino-acids (Kopp *et al*., 2013) necessary for protein synthesis and algal filament growth. However, the δ^15^N values of most of the decalcified endolithic strains were higher than those of the residual skeletal organic matrix after carbonate dissolution (Fig. 4; not an effect of the acid-treatment used for decalcification, see results). Thus, the skeletal organic matrix, known to be rich in amino-acids and glycoproteins (Marin *et al*., 2016) could be an additional heterotrophic nitrogen source for *Ostreobium* endoliths. Other microbial processes might be at play, and contribute to nitrogen isotope fractionation, linked to N_2_ fixation, nitrate reduction, and cycling activities of associated endolithic bacterial/fungal microorganisms in the coral skeleton microbiome (Ferrer & Szmant, 1988; Wegley *et al*., 2007; our results showing the presence of fungal and bacterial fatty acid markers in cultures of *Ostreobium*). The demonstrated uptake of nitrate by *Ostreobium* strains, especially high for the free-living phenotype but also significant for the endolithic phenotype, highlights an important role of *Ostreobium* in reef nitrogen cycling, as suggested in early works on nutrient regeneration in living corals (Risk & Müller, 1983; Ferrer & Szmant, 1988).

### Conceptual model of C and N sources for Ostreobium

In light of our results, we propose a model of nutritional sources for *Ostreobium* algae, illustrated in Figure 6. In this conceptual model, an endolithic filament emerges from the coral carbonate surface as an epilith, which can then be detached and grown in seawater as a free-living filament. Dissolved CO_2_ and bicarbonate ions (HCO_3_^-^) from seawater are in direct contact with epilithic *Ostreobium* filaments, and diffuse from gallery opening to the interstitial space between endolithic filaments and skeleton. Dissolved CO_2_ is also provided by respiration. Bicarbonate ions are converted to CO_2_ via the activity of carbonic anhydrase enzymes (CA) (Shashar & Stambler, 1992). Thus, CO_2_ (from seawater or formed after conversion of HCO_3_^-^ by the CA) diffuse inside the *Ostreobium* filament and is fixed in chloroplasts by the ribulose-1,5-bisphosphate Carboxylase/Oxygenase (RuBisCO) enzyme for photosynthetic production of C-rich organic compounds (i.e. glucose, fatty acids) used for growth. At the dissolution front of endolithic *Ostreobium*, inorganic C might be taken up directly by the filament for photosynthesis, or after conversion of organic C from dissolved organic matter (DOM) released by the breakdown of the skeletal organic matrix and recycled by associated endolithic bacteria and fungi. Dissolved inorganic nitrogen (DIN, in this study NO_3_-) is provided by the incubation seawater and diffuses to the interstitial space between endolithic filament and coral skeleton. Assimilation of DIN may be combined to organic N from other microbial processes (hypothesized by Risk & Müller, 1983; Ferrer & Szmant, 1988) and the mobilization of DOM from biogenic coral carbonate dissolution. The exact metabolic processes of organic and inorganic N acquisition, i.e. their diffusion or active transport into *Ostreobium* filaments, remain to be investigated.

**Figure 6:**
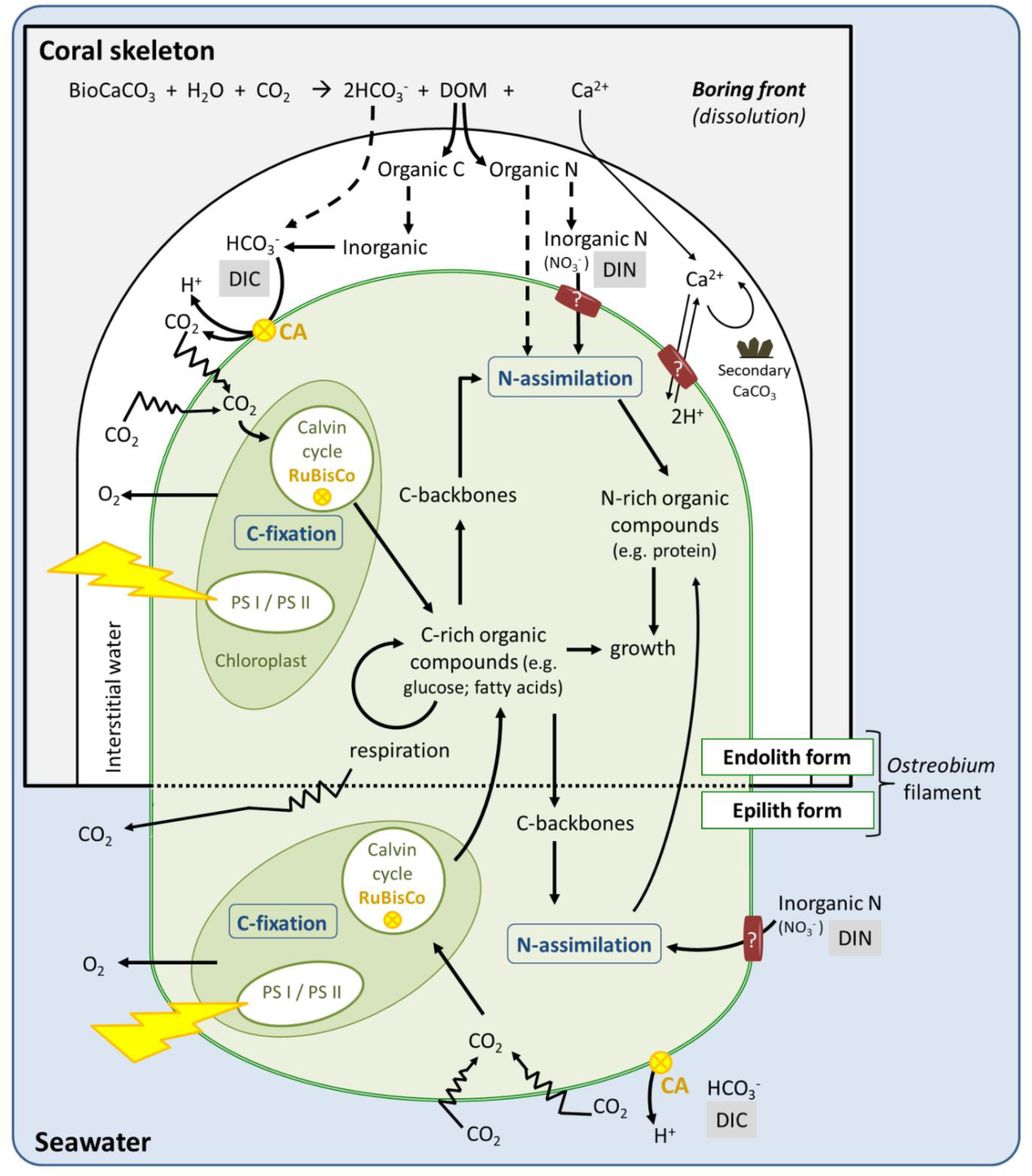
Conceptual model of C and N sources for *Ostreobium* filaments as endolithic and epilithic/free-living growth form. BioCaCO_3_: biogenic carbonate. ?: putative ion transporter. Dotted line indicates hypothetical use of DIC from carbonate dissolution, and organic N by endoliths.

We suggest that endolithic *Ostreobium* may have mixed inorganic and organic C and N sources, compared to free-living forms which fix inorganic C and assimilate N through photosynthesis-dependent processes. This nutritional plasticity hypothesis proposes that *Ostreobium* filaments may change their metabolism to adapt to their habitat and/or environmental conditions (i.e. both seawater chemistry and thickness/composition of associated living coral tissues or epilithic biofilms), shifting from photoautrophy in epilithic/free-living growth form to mixotrophy in endolithic growth form. This could explain the large bathymetrical distribution of endolithic *Ostreobium*, including depths below the photic zone (Vogel *et al*., 2000), and the rapid colonization of freshly killed corals by initially present endolithic *Ostreobium* filaments (Leggat *et al*., 2019). Genotype-driven differences in C and N acquisition strategies should be further investigated, in light of our results showing highly variable δ^13^C and δ^15^N values and DIC and DIN uptake rates among strains within and between genetic lineages. Further isolation and cultivation of strains from diverse genetic lineages under diverse nutrient conditions will definitely help to better understand *Ostreobium* eco-physiology and the responses of microboring communities to global changes (e.g. Carreiro-Silva *et al*., 2005 for eutrophication effects, and Tribollet *et al*., 2009 for ocean acidification effects).

### Conclusion

This study highlights habitat-driven changes in chlorophylls, fatty acids and stable isotope values (δ^13^C, δ^15^N) of coral-associated *Ostreobium* strains from two genetic lineages, providing novel information on their nutritional value as free-living/epilithic and endolithic filaments. Lower photosynthetic assimilation and nitrate uptake from seawater by endolithic *versus* free-living phenotypes may be combined for some *Ostreobium* strains with heterotrophic mobilization of organic matter from microbial associates and recycled from the skeletal organic matrix after active dissolution of the carbonate biomineral. We propose that nutritional plasticity may depend on habitat and environmental conditions, with *Ostreobium* filaments shifting from photoautotrophic to mixotrophic lifestyle when free-living filaments (or propagules in seawater) colonize reef carbonates as endoliths. The dual isotope tracer approach used here opens the way to further study the biogeochemical cycling and trophic ecology of these cryptic algae inhabiting coral holobionts and reef carbonates, to help understand coral reef resilience to global changes.

## Materials and Methods

### Isolation of *Ostreobium* strains

Strains of *Ostreobium* were isolated from healthy *Pocillopora acuta* Lamarck 1816 coral colonies (genetically identified with mtORF markers after Johnston *et al*., 2017, also called *Pocillopora damicornis* type *beta* coral), growing in closed-circuit at the Aquarium Tropical du Palais de la Porte Dorée (Paris, France originally from Indonesia). One apical branch fragment (‘apex’, ∼10 mm length) was sampled from each of three distinct *P. acuta* colonies. Skeleton was cleaned off coral tissues containing *Symbiodinium* and other microbes using a blast of pressurized filtered (0.2 µm) seawater (WaterPik^®^ method; Johannes & Wiebe, 1970), then crushed into big pieces using autoclaved mortar and pestle. Fragments were separated in 3 fractions, each deposited in an individual microplate well filled with ∼5 mL of Provasoli Enriched Seawater medium (PES; Provasoli, 1968) supplemented with penicillin G sodium (100 U mL^-1^) and streptomycin sulfate (100 µg mL^-1^). Cultures were incubated at 25°C with 30-40 rpm orbital shaking and a 12h light / 12h dark cycle of illumination with white fluorescent light tubes at photosynthetic photon flux density of 31±5.5 μmol.m^-2^.s^-1^ (measured with a spherical quantum sensor Li-Cor, USA). Culture medium was changed every 3-4 weeks. After ∼1 month, epilithic filaments emerged from the skeletal carbonate chips, identified as *Ostreobium* by their typical morphology in inverted light microscopy (Olympus CK40-SLP). Other green and red algae could sometimes be observed. Epilithic *Ostreobium* filaments were pulled out or cut with a scalpel from the skeleton surface and then serially transferred to fresh PES medium for propagation into monoalgal cultures of free-living filaments. The obtained *Ostreobium* cultures were not axenic as they also contained bacteria and sometimes dinoflagellates, which population densities were controlled by regular subculturing (passage) of isolated algal filaments. For each strain, the number of successive subcultures since initial sampling from skeletal piece was recorded. Endolithic cultures were obtained from the free-living cultures, by colonization during several months of pre-bleached native coral carbonate chip substrates (see below). Strain vouchers have been deposited in the Collection of microalgae and cyanobacteria of the Muséum national d’Histoire naturelle in Paris, France (Table 1).

### Taxonomic assignation to *rbc*L clade (phylotype)

For each algal strain, free-living filaments were subsampled for DNA extraction (with DNeasy PowerSoil^™^ Kit, Qiagen Laboratories Inc., CA) and genotyping based on sequences of the chloroplast-encoded *rbc*L gene marker coding for the large subunit of the ribulose-1,5-bisphosphate carboxylase -RuBisCo-enzyme, according to our previous classification scheme (Massé *et al*., 2018). A partial fragment (∼1134 nt) of the 1428 nt chloroplast-encoded *Ostreobium rbc*L gene was amplified with the following oligonucleotide primer pair, specific to the Bryopsidales order within the Ulvophyceae: *rbc*LF250 [5’ GATATTGARCCTGTTGTTG GTGAAGA 3’] modified from Gutner-Hoch & Fine (2011) and *rbc*L1391R [5’TCTTTCCA AACTTCACAAGC 3’] (Verbruggen *et al*., 2009). A smaller ∼390-440 nt fragment of the *rbc*L gene was obtained for two cultures which DNA was more difficult to amplify, using the primer pair: *rbc*LF250 and *rbc*LR670 [5’ CCAGTTTCAGCTTGWGCTTTATAAA 3’] modified from Gutner-Hoch & Fine (2011). Amplification reactions were performed in 25 µL volume containing 1 µL DNA extract template, 0.5 µL of each primer (10 µM final concentration), 2 µL MgCl_2_ (25 mM), 0.5 µL dNTP (10 mM), 5 µL GoTaq 5X Buffer, 0.125 µL enzyme GoTaq® G2 Flexi DNA Polymerase (Promega, France) in sterile water. Cycling conditions were 4 min at 94°C, 40 cycles of [30 s at 94°C, 45 s at 55°C, 90 s at 72°C], and 5 min terminal extension at 72°C. Amplified fragments were visualized on 1 % agarose gel with SYBRGold or SYBRSafe (Invitrogen). Purified *rbc*L amplicons (NucleoSpin® gel and PCR clean-up kit, Macherey-Nagel, France) were either Sanger-sequenced directly or cloned into pGEM-T easy vector plasmids using competent *Escherichia coli* JM109 cells (Invitrogen, France). Plasmid DNA of insert-containing colonies was extracted (Wizard Plus SV Minipreps, Promega, France) and Sanger-sequenced in both directions by Eurofins Genomics (Germany). Forward and reverse sequences were assembled and checked manually with BioEdit software. Taxonomic assignation of *Ostreobium* strains was determined by BLASTn against GenBank database. All sequences generated during this study have been deposited in Genbank under accession numbers MK095212 to MK095220 (Table 1).

### Bio-erosive potential in culture conditions

Pieces of *Pocillopora acuta* coral skeletons (bared of coral tissue) were bleached by immersion during one to three days at room temperature in sodium hypochlorite (NaClO 48%, commercial bleach) to remove potential residual algae, coral tissues and surface organic matter. They were rinsed thoroughly with tap water then deionized water and finally 70% ethanol then air-dried under laminar flow hood. For each *Ostreobium* strain, bleached skeletons were put in contact with free-living filaments to test whether these filaments kept their erosive activity and would be able to colonize their native carbonate substrate, as indicated by (i) filament attachment to skeletal surfaces and (ii) progressive color change of the white, bleached skeletal chip to green *Ostreobium*-invaded chip. These visible signs of colonization were confirmed microscopically : a bleached coral skeleton exposed for about 3 months to filaments of *Ostreobium* 010 was fixed in 4% paraformaldehyde in sucrose containing phosphate-buffer, dehydrated in ethanol series (50%, 70%, 96%, 100%), gradually infiltrated with resin (1:2; 1:1; 3:1; v:v ethanol-Spürr resin) then resin-embedded (100% Spürr). Sections cut with a circular diamond saw were polished down to ∼20-30 µm, slightly decalcified with formic acid 5% and rinsed with deionized water. These thin sections were either stained with 5% toluidine blue and coverslipped in Spürr for light microscopy observations (Olympus CK40-SLP) or gold-coated and mounted for scanning electron microscopy observations (Hitachi SU 3500, MNHN PtME platform for electron microscopy) of endolithic filaments.

Thus, for each free-living *Ostreobium* strain, we also obtained its corresponding endolithic growth form. Before all biochemical and physiological analyses, coral skeletal chips colonized by endolithic *Ostreobium* were scraped with a sterile toothbrush, or blasted with a jet of pressurized filtered seawater (Waterpik^®^ method) and transferred to 0.2µm filtered seawater for 24h to 3 days, in order to remove most outgrowing epilithic filaments and select the targeted microboring endolithic growth form.

### Photosynthetic and accessory pigment analysis

Free-living *Ostreobium* isolates (5 strains) and their corresponding endolithic growth forms (3 strains, due to limited available biomass) belonging to lineage P1 (see Table 1, less frequent strains P12/P14 were not studied) were washed three times with large volumes (1:25) of autoclaved deionized water to mechanically remove most surface bacteria and dinoflagellate contaminants and decrease the salt concentration before organic extractions. To evaluate variability in pigment profile across subcultures within same strain, two successive passages (i.e. subcultures) of 3 free-living *Ostreobium* strains were analyzed (see Table 1). Furthermore, bleached *Pocillopora acuta* coral skeletons (uncolonized substrate, n=3) were analyzed as controls for potential residual pigments. The samples were lyophilized and ground to powder. Final dry weights were of 20.6±12.2 mg for free-living forms, 2253±1058 mg for endolithic forms (without carbonate dissolution), and 1256±361 mg for control bleached coral skeletons (without carbonate dissolution). Samples were extracted twice in dichloromethane/methanol (CH_2_Cl_2_/MeOH; 1:1 v:v), sonicated (10-15 minutes) in ice in the dark to prevent pigment degradation, and then filtered. After evaporation, organic extracts were re-solubilized in a mixture of CH_2_Cl_2_/MeOH (1:1 v:v) to a final concentration of 10 mg mL^-1^. Using high-performance liquid chromatography (HPLC, Agilent 1220 infinity) equipped with a diode array detector (DAD), each extract was analyzed on a reverse phase column (C18 Capcell Pak, Shiseido, 4.6 x 250 mm). Elution solvents were those used by Frigaard *et al*. (1996), i.e. solvent A (methanol:acetonitrile:water, 42:33:25) and solvent B (methanol:acetonitrile:ethyl acetate, 39:31:30) starting with the gradient elution of solvent B from 40 to 100% during 60 minutes (flow 1 mL min^-1^). The wavelengths of 664 nm and 470 nm were used to visualize chlorophyll *a* and its allomers, as well as other chlorophylls/carotenoids, respectively. All peaks were characterized by their retention time (RT) and maxima absorption spectra. The peaks corresponding to chlorophyll *a* (chl *a*) and *b* (chl *b*) were identified (RT and maxima absorption spectra) by direct injection of chl *a* and chl *b* standards (Sigma-Aldrich C-5753 and 00538, respectively) under the same HPLC conditions. Moreover, one *Ostreobium* free-living strain extract was supplemented with both standards as internal controls in order to confirm their respective localization within the HPLC profiles (Fig. S2a). Calibration curves were established for chl *a* and chl *b* standards to quantify chl *a* and chl *b* within the different HPLC profiles of *Ostreobium* cultures (relative proportion and chl *b* : chl *a* ratio).

### Fatty acid analysis

Endolithic and free-living *Ostreobium* strains (total 8 strains, from P1 and P12/P14, see Table 1), as well as control bleached skeletons (n=4), were lyophilized and ground rapidly (a few seconds) to powder. Two to three subcultures were analyzed for each of 1 endolithic and 3 free-living *Ostreobium* strains, to evaluate variability in fatty acid composition across subcultures within same strain. Final dry weights were 709±378 mg for endolithic forms (without carbonate dissolution), 2.44±0.3 mg for free-living forms, and 504±100 mg for control skeletons (without carbonate dissolution). Crushed samples were frozen at −20°C until lipid extraction, according to a method modified from Bligh & Dyer (1959). Briefly, after sonication for 20 min in water/methanol/chloroform (H_2_O/MeOH/CHCl_3;_ 1:2:1, v:v:v), H_2_O/CHCl_3_ (1:1 v:v) was added to form an aqueous-organic two-layer system. The lipid-containing lower chloroform phase was recovered after centrifugation (3000 rpm, 5 min). The aqueous phase was re-extracted a second time. Combined chloroform phases were evaporated under a nitrogen stream. Saponification of extracts was performed at 90°C for 90 min with a solution of NaOH(2M):MeOH (1:2, v:v), followed by acidification with ultra-pure HCl solution (35%). Lipids were recovered by centrifugation (3000 rpm, 5 min) after adding chloroform (2 x 1.5 mL). Chloroform phases were again combined and evaporated under a nitrogen stream. Transmethylation of total lipids was conducted using Boron trifluoride-methanol (BF_3_) at 90°C for 10 min. Lipids were finally re-extracted and washed in H_2_O:CHCl_3_ (1:1, v:v). Chloroform phases recovered were evaporated under a nitrogen stream before being solubilized in hexane and stored at −20°C for analysis by gas chromatography (Varian 450-GC and VF-WAXms column 30 m x 0.25 mm; 0.25 μm film thickness). Peaks of fatty acids were identified using GC-mass spectrometry (Varian 220-MS) and comparison with retention times of commercial standards (Supelco 37, Supelco PUFA N°1 and Supelco PUFA N°3).

### Stable isotope values and measurements of photosynthetic C, N assimilation

Isotope dual labeling experiments were conducted for endolithic and their free-living *Ostreobium* counterparts (n=4 pairs, 2 per each genetic lineage P1 and P12/P14; see Table 1). Skeletons colonized by endolithic forms were brushed to remove epilithic filaments as described above, and all culture samples (free-living and endolithic growth forms) were transferred for 3 days before the labeling experiment in Artificial SeaWater (ASW-adapted from Harrison *et al*., 1980) in order to rinse the nutrient rich PES medium. The isotope labeling pulse was conducted during 8h in white light at 30.5±2.5 μmol photons.m^-2^.s^-1^ Photosynthetically Active Radiation (PAR) measured with a LI-250A (LI-COR) quantum-meter equipped with a submersible spherical Micro Quantum Sensor US-SQS/L (WALZ) or in control darkness (inside aluminium foil-wrapped box), starting at the beginning of the light period. Samples of *Ostreobium* cultures (n=4 pairs) or control bleached skeletons (uncolonized, n=3) were incubated at 25°C, under gentle orbital shaking (30-40 rpm, Infors Minitron incubator, Fr) in glass jars with plastic lids, filled to ¾ of their volume with 25 mL of ASW (initially free of bicarbonate and nitrate) supplemented with 2 mM ^13^C-bicarbonate (NaH^13^CO_3_, 99 atom% [Sigma-Aldrich]) and 5 µM ^15^N-nitrate (K^15^NO_3_, 98 atom%[Sigma-Aldrich]) or in ‘control unlabeled ASW’ supplemented with 2 mM natural abundance sodium bicarbonate [Sigma-Aldrich] and 5 µM natural abundance potassium nitrate [Sigma-Aldrich]. The labeling experiment was repeated later in a separate experiment for an additional strain 010 (lineage P1), to increase strain and technical replication under similar light conditions.

Seawater pH was measured in glass jar with samples at beginning and end of the 8h experiment (NBS scale; Electrode pH Mettler Toledo Inlab 413 with a pHmeter Mettler Toledo MP220). Control bleached skeletons (uncolonized by endoliths) showed a slight decrease of pH (−0.04) at the end of the labeling pulse in light conditions. Assuming a similar trend in dark conditions, we added this correction value from those pH values obtained for all endolithic strains.

At the end of the 8h labeling pulse, samples were rinsed three times with ASW (free of bicarbonate and nitrate). Endolithic *Ostreobium* and control bleached skeletons were decalcified with formic acid 5% and rinsed with deionized water. To assess the effect of the acid-decalcification treatment on ^13^C and ^15^N isotope enrichments (labeling intensities), strain 010 labeled in free-living form was divided in two parts: one part was lyophilized directly and the other part was treated with formic acid 5%, similar to the carbonate removal protocol to analyze decalcified endolithic forms and control skeletons. The effect of the acid-decalcification treatment was also tested on δ^13^C and δ^15^N stable isotope values of free-living strain 010 untreated or treated with formic acid 5%.

To increase replication of measurements of C and N stable isotope values across subcultures, strains, and genetic lineages, additional samples were analyzed, of endolithic *Ostreobium* strains (not decalcified, or decalcified with formic acid 5%) and their corresponding free-living forms, and control bleached skeleton substrate (see Table1).

All organic samples were lyophilized, weighted (0.5-2 mg) into tin capsules and sent for bulk isotope dual ^13^C and ^15^N analyses on an elemental analyzer interfaced to a continuous flow isotope ratio mass spectrometer (EA-IRMS) at UC Davis Stable Isotope Facility (California, USA) using bovine liver, glutamic acid, and ^15^N-enriched alanine as internal standards. Alternately, the series of *Ostreobium* 010 samples and additional unlabeled replicate strains were analyzed in an automated combustion system (EA Flash 2000 Thermo) interfaced with a DeltaV Advantage Thermo continuous flow IRMS at the MNHN SSMIM facility (Paris, France) using normal abundance alanine (0.3 mg) as internal standard. The analytical uncertainties within the SSMIM run estimated from 14 repeated analyses of the alanine laboratory standard was lower than 0.13 ‰ (1SD) for δ15N values and lower than 0.10 ‰ (1SD) for δ13C values.

The Carbon to Nitrogen (C/N) ratio was calculated for each sample following the equation: C/N ratio = (C Amount/12.01) / (N Amount/14.006)

Natural isotope abundances were expressed according to the delta notation:

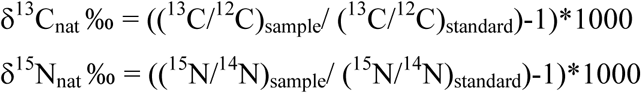

with R_standard_ = 0.01123 for ^13^C/^12^C (Vienna Peedee Belemnite calcite standard) and 0.00367 for ^15^N/^14^N (atmospheric N_2_ standard).

For the carbonate samples, i.e. the undecalcified coral carbonate substrate controls (uncolonized) or endolithic growth forms (coral skeletal chips colonized by *Ostreobium)*, an alternative sample preparation method was used, the Kiel IV (Thermo) automated carbonate preparation device interfaced to a Delta V Advantage IR-MS (Thermo), at the MNHN SSMIM facility. Sample powders (40-60 µg) were analyzed individually. The δ^13^C values were expressed versus the Vienna-Pee Dee Belemnite calcite standard, and corrected by comparison with a laboratory standard (*Marbre LM*) normalized to the NBS 19 international standard (δ^18^O values are not presented here). The mean analytical precision within the run was calculated from 8 measurements of the *Marbre LM* averaging 0.038 ‰ (1SD) for δ^13^C values.

For isotopically labeled samples (free-living and decalcified endolithic *Ostreobium*, and control decalcified skeletons, i.e. skeletal organic matrix), all ^13^C and ^15^N-enrichment levels were expressed according to the delta notation: δX (‰) = ((R_sample_/R_control_)-1)*1000. R_sample_ is the ratio ^13^C/^12^C or ^15^N/^14^N of labelled samples, and R_control_ is the measured ratio of corresponding control non-labeled samples (i.e. natural isotope abundance ratio).

Labeling values for decalcified endolithic forms were corrected by subtracting the average of 3 replicate values obtained for control decalcified skeletons, corresponding to non-specific adsorption of the isotopic label to the skeletal organic matrix (means of 0.4±2 ‰ for enriched δ^13^C and 0.9±0.2 ‰ for enriched δ^15^N). We also tested the effect of formic acid treatment and showed that it introduced variability but did not change average ^13^C and ^15^N enrichments in free-living *Ostreobium* (^13^C-enrichment of 585±106 *versus* 553±19.7 ‰ and ^15^N-enrichment of 384±222 *versus* 377±42 ‰, with or without formic acid respectively; n=3 technical sub-replicates of strain 010). We thus assumed that the acid-formic treatment did not significantly affect enrichment results obtained for decalcified endolithic *Ostreobium*.

### Data analysis

Multivariate analysis of pigment compositions was performed using PRIMER 5 software. A triangular similarity matrix was created using Bray-Curtis similarity coefficient, followed by non-metric multidimensional scaling (nMDS) to spatially visualize samples of endolithic, free-living *Ostreobium* strains and control skeletons with similar composition. Similarity and dissimilarity percentages obtained by SIMPER analysis allowed to determine which pigments (or RT of peaks in case of non-identified pigment) drove the observed differences between datasets (i.e. endolithic *versus* free-living *Ostreobium versus* control skeletons).

Principal component analysis (PCA) was performed on % level of fatty acids using R software (version 3.6.2), to reveal spatial variability among strains, genetic lineage, and growth forms of *Ostreobium*, and identify FAs that explain most the variance in our datasets.

For δ^13^C and δ^15^N stable isotope values, parametric Student-test were performed with R software (version 3.6.2) to determine if the genetic lineage (P1 *versus* P12/P14) influenced δ^13^C and δ^15^N values for both *Ostreobium* growth form. Then, an analysis of variances (ANOVA) was performed on δ^13^C and δ^15^N datasets, with post hoc Student-Newman-Keuls (SNK) tests, highlighted differences between *Ostreobium* growth forms and with their habitat (coral skeleton substrate). Significance threshold were set to p-value <0.05.

## Supporting information

Supplementary Informations

## Acknowledgments

This work was supported by UPMC IPV (‘Interface Pour le Vivant’) doctoral grant to AM, MNHN grants ATM 2017-2018 ‘ECTOSYMBIOCORAL’ and Aviv 2017 ‘Photosymbioses marines’ to IDC. Additional funding support came from ISPL-LOCEAN lab UMR7189 to AT and MCAM lab UMR7245 to IDC. We thank the director and staff of the Aquarium Tropical, Palais de la Porte Dorée (Paris, France) for providing *Pocillopora acuta* colonies from which *Ostreobium* strains were isolated. We thank Denis Fiorillo at the Service de Spectrométrie de Masse Isotopique of the Muséum national d’Histoire naturelle (SSMIM, MNHN Paris, France) for help with organic and carbonate C and N stable isotope analyses. We also thank Pr. Karim Benzerara (IMPMC lab UMR7590) for discussions, and the editor and five anonymous reviewers who helped improving the manuscript.

## Authors Contributions

A.M, I.D.C, and A.T conceived the research and designed the experimental protocols, A.M performed the experiments with help from C.Y. for strain isolation, C.S. for stable isotope experiments, A.L. for pigment extraction and N.T. for fatty acid analyses. T.M, M-L.B-K and A.C. helped analyze and discuss the data. A.M., A.T. and I.D-C. wrote the manuscript, which was commented by all authors. A.M. A.T. and I.D.C. revised the manuscript.

