## Supplementary Informations for "Functional diversity of microboring *Ostreobium* algae isolated from corals"

**Running head**: Physiology of an algal microborer of coral carbonate

**Supplementary Informations**

**Figure S1**: ***rbc*L phylogeny of *Ostreobium* sp. strains isolated from coral host *Pocillopora acuta***. Maximum likelihood tree based on *rbc*L (161 nt) sequence analysis: cloned OTUs (>99% similarity) were aligned to chloroplast genomes from reference *Ostreobium* strains and cloned *rbc*L fragments from aquaria and reef corals (Massé *et al*., 2018, and pers. com.), with one Ulvaceae (Ulvale) and one Caulerpaceae (Bryopsidale) as outgroup in the class Ulvophyceae (1000 bootstraps). Clade P1 forms a species-level entity (OTU>99%) whereas clades P12 and P14 are clustered into a more phylogenetically distant genus-level entity (OTU>97%).

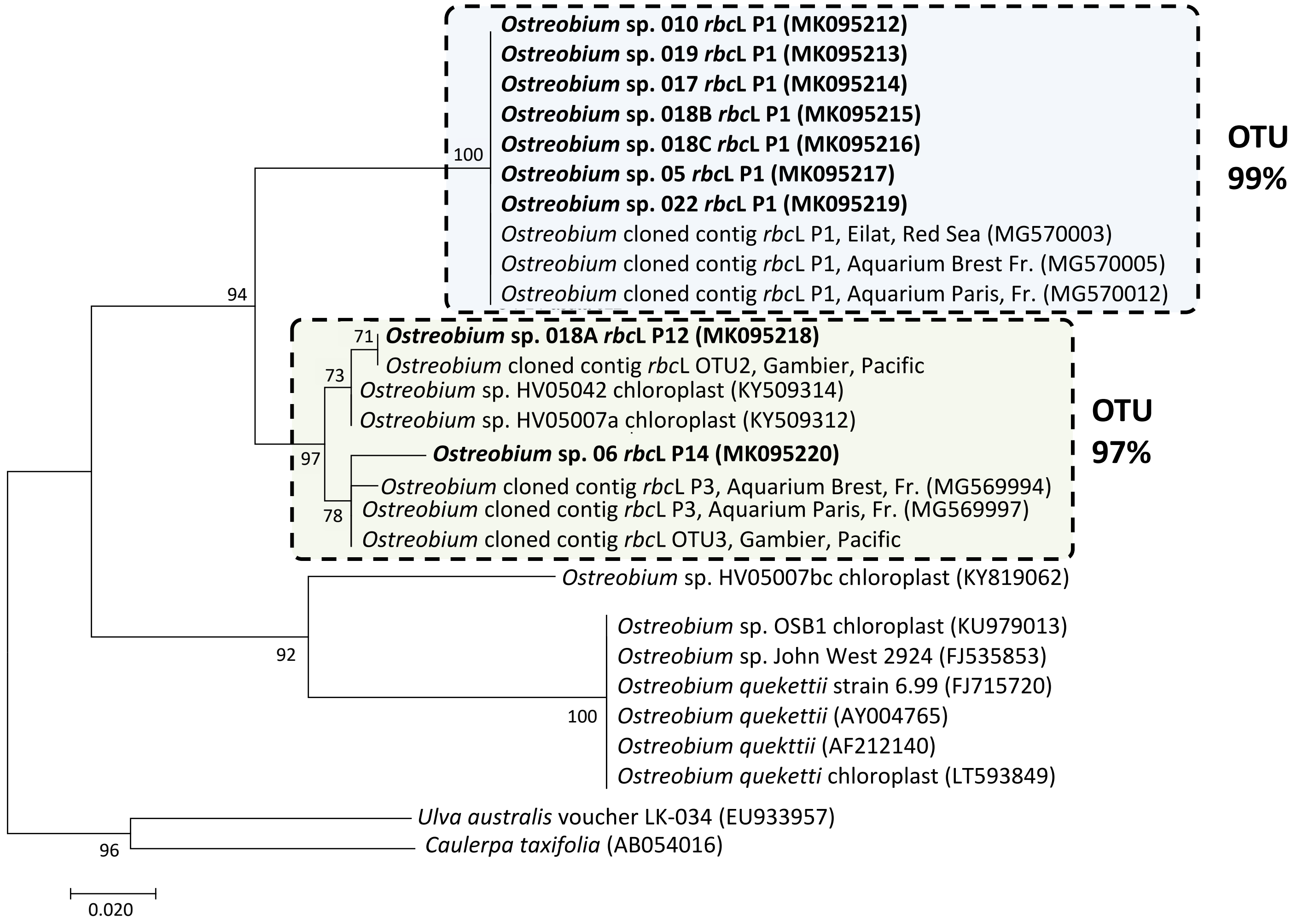

**Figure S2: Chromatographic profiles (HPLC) of the pigment composition from *Ostreobium* strains.** (a) HPLC profiles of a representative free-living *Ostreobium* strain (clade P1, 018B), alone or supplemented with chlorophyll *a* and *b* standards at 664 nm and 470 nm. (b) Overlayed HPLC profiles from a representative *Ostreobium* strain (clade P1, 019) in endolithic *versus* free-living growth form, at 470 nm and 664 nm, with chlorophyll *a* and *b* identified. The relative proportion of each pigment has been reduced to a percentage scale.

**a**

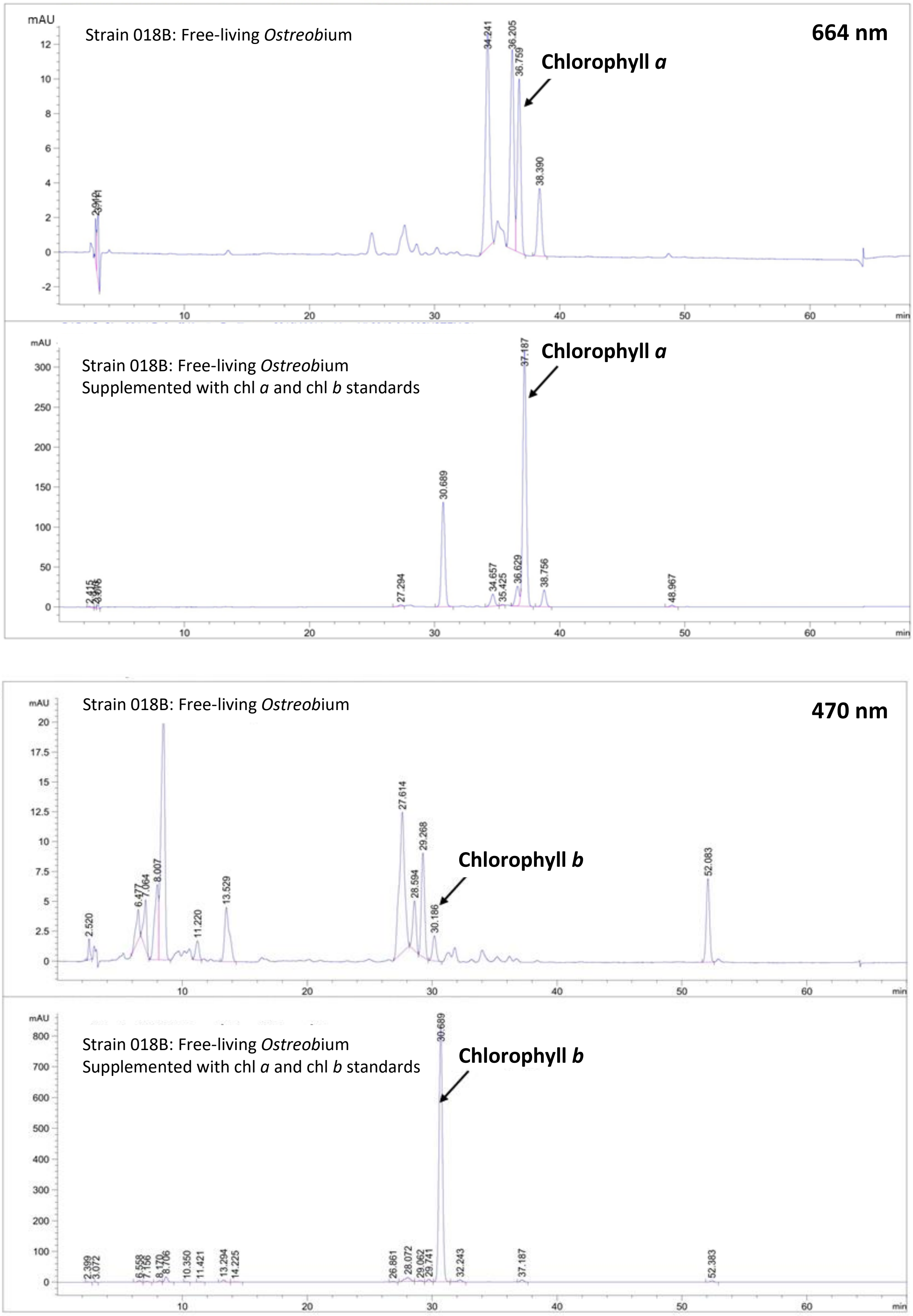

**b**

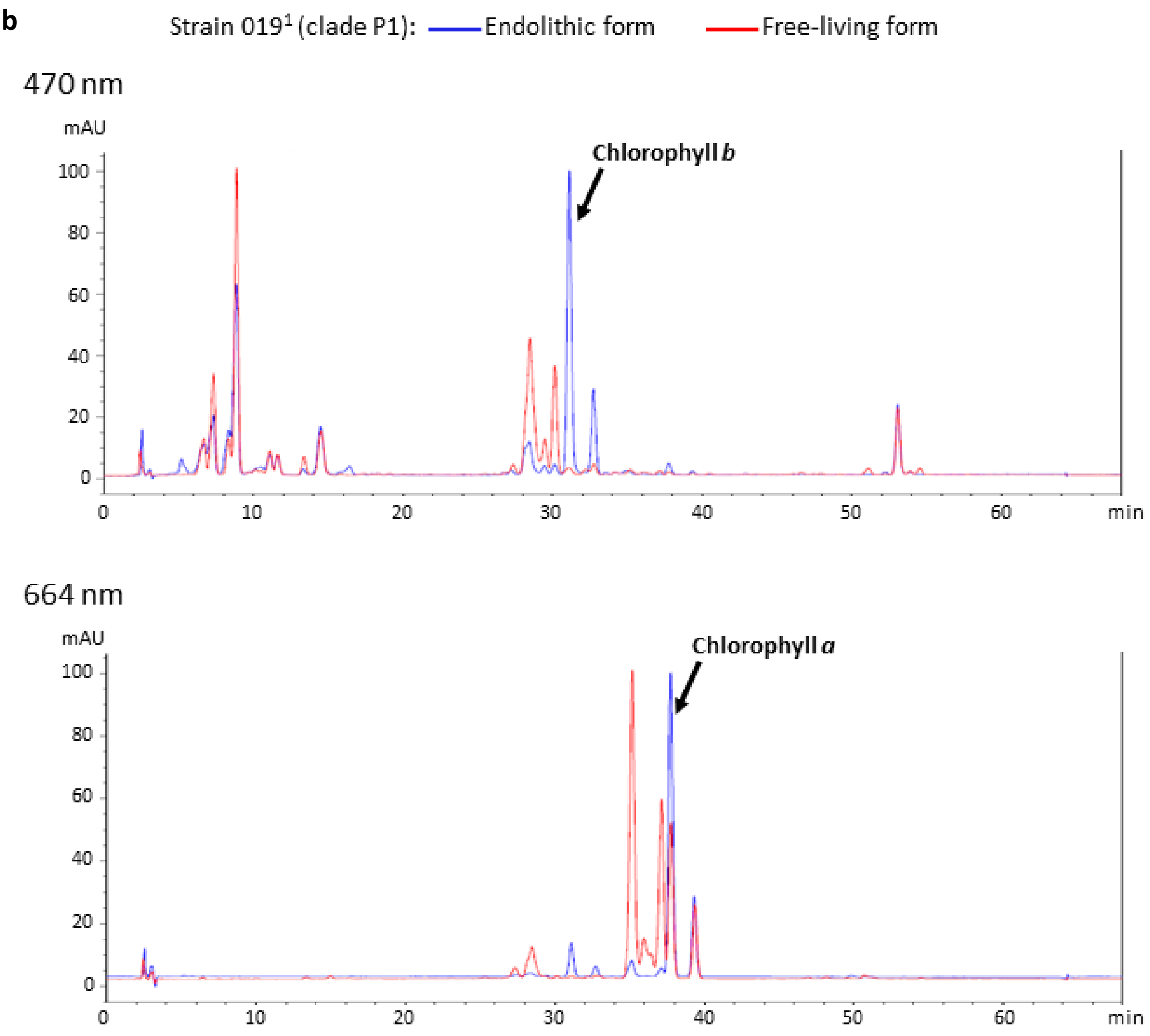

Strain 019[1] (clade P1):
Endolithic form
Free-living form
470 nm
mAU
Chlorophyll *b*
100
80
60
40
20
0
0
10
20
30
40
50
60
min
664 nm
mAU
Chlorophyll *a*
100
80
60
40
20
0
0
10
20
30
40
50
60
min

**Figure S3: Chromatographic profiles (GC) of fatty acids composition of (a) a free-living *Ostreobium* strain compared to (b) its endolithic growth form.**

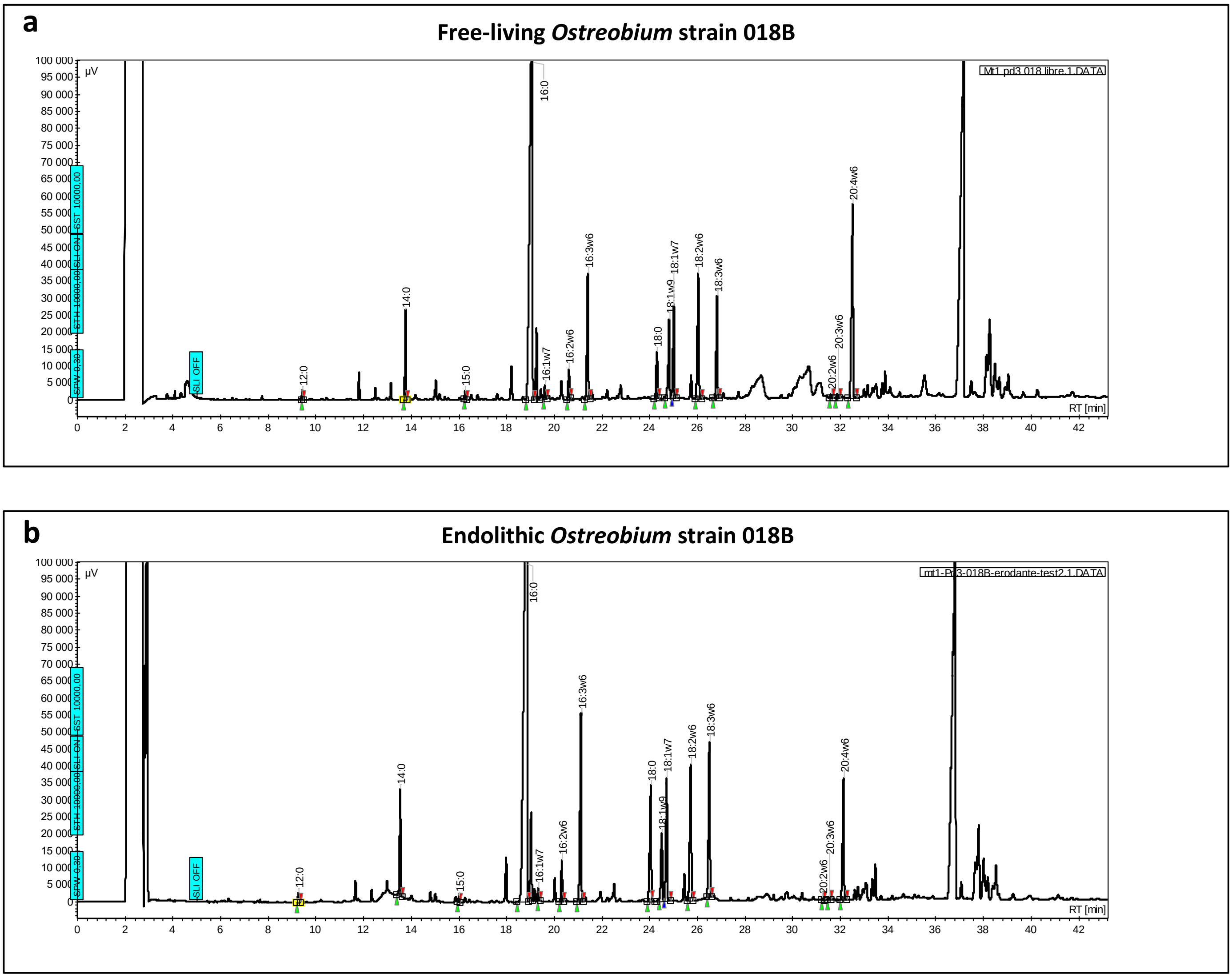

**Table S1: Pigment composition in endolithic *versus* free-living *Ostreobium* (clade P1), as well as in control bleached coral carbonate skeletons.** Mean values of the percentage of total pigment content measured in HPLC profiles ± Standard Error, SE. (number of extracts with detected peak/ total number of extracts for each growth form).

| Mean retention time ± SD (min) | Identified peak | Maxima Absorption (nm) | Free-living *Ostreobium* (% of total pigments: mean±SE) | Endolithic *Ostreobium* (% of total pigments: mean±SE) | |
|---|---|---|---|---|---|
| | | | Clade P1 (n=8) | Clade P1 (n=3) | Control bleached skeletons (n=3) |
| 2.1 | | 428, 652 | 0.04±0.04 (1/8) | 0±0 (0/3) | 0±0 (0/3) |
| 2.3 | | 400, 668 | 0.04±0.04 (1/8) | 0±0 (0/3) | 0±0 (0/3) |
| 2.5 | | 428, 452 | 0.37±0.37 (1/8) | 0±0 (0/3) | 0±0 (0/3) |
| 2.6±0.1 | | 444, 632 | 1.07±0.63 (3/8) | 2.36±1.54 (3/3) | 0±0 (0/3) |
| 2.9 | | 444, 630 | 0.22±0.22 (1/8) | 0±0 (0/3) | 0±0 (0/3) |
| 2.9 | | 440, 664 | 0±0 (0/8) | 0.30±0.30 (1/3) | 0±0 (0/3) |
| 2.9 | | 464 | 0.17±0.17 (1/8) | 0±0 (0/3) | 0±0 (0/3) |
| 5.1 | | 442 | 0.22±0.22 (1/8) | 0±0 (0/3) | 0±0 (0/3) |
| 5.2 | | 434 | 0±0 (0/8) | 0.30±0.30 (1/3) | 18.23±11.42 (2/3) |
| 5.4±0.1 | | 468 | 0.29±0.24 (2/8) | 0±0 (0/3) | 0±0 (0/3) |
| 5.4±0.2 | | 472 | 2.0±1.32 (2/8) | 0.35±0.35 (1/3) | 0±0 (0/3) |
| 6.6±0.2 | | 444, 468 | 2.21±0.78 (7/8) | 6.50±3.96 (3/3) | 0±0 (0/3) |
| 6.6 | | 448, 468 | 0.43±0.43 (1/8) | 0±0 (0/3) | 0±0 (0/3) |
| 7.2±0.2 | | 416, 436, 468 | 2.95±0.66 (8/8) | 4.18±0.74 (3/3) | 0±0 (0/3) |
| 8.2±0.2 | | 416, 440, 472 | 2.34±0.81 (5/8) | 3.72±1.29 (3/3) | 0±0 (0/3) |
| 8.7±0.2 | | 448, 468 | 16.56±1.40 (8/8) | 13.13±1.23 (3/3) | 0±0 (0/3) |
| 9.0 | | 436 | 0±0 (0/8) | 0±0 (0/3) | 16.32±11.90 (2/3) |
| 9.9±0.3 | | 448, 476 | 1.15±0.59 (4/8) | 0.36±0.14 (3/3) | 0±0 (0/3) |
| 10.1 | | 452, 476, 648 | 0.02±0.02 (1/8) | 0±0 (0/3) | 0±0 (0/3) |
| 10.4±0.2 | | 440, 464 | 0.52±0.23 (5/8) | 0.15±0.02 (3/3) | 0±0 (0/3) |
| 10.8 | | 440, 472, 660 | 0.02±0.02 (1/8) | 0±0 (0/3) | 0±0 (0/3) |
| 10.9±0.2 | | 448, 476 | 0.68±0.25 (5/8) | 0.66±0.33 (2/3) | 0±0 (0/3) |
| 11.4±0.3 | | 332, 436, 460 | 1.08±0.11 (8/8) | 0.81±0.11 (3/3) | 0±0 (0/3) |
| 12.3 | | 450, 628 | 0.1±0.08 (2/8) | 0±0 (0/3) | 0±0 (0/3) |
| 13.4±0.6 | | 452, 632 | 3.83±0.79 (8/8) | 0.41±0.07 (3/3) | 0±0 (0/3) |
| 14.2±0.4 | | 448, 476 | 1.01±0.52 (3/8) | 2.44±0.80 (3/3) | 0±0 (0/3) |
| 15.8 | | 452, 644 | 0.01±0.01 (1/8) | 0±0 (0/3) | 0±0 (0/3) |
| 16.3±0.1 | | 436 | 0.04±0.04 (1/8) | 0.17±0.17 (1/3) | 65.45±9.92 (3/3) |
| 16.8±0.3 | | 452, 636 | 0.22±0.09 (4/8) | 0±0 (0/3) | 0±0 (0/3) |
| 17.2±0.5 | | 416, 636 | 0.03±0.03 (3/8) | 0±0 (0/3) | 0±0 (0/3) |

| | | | | | |
|---|---|---|---|---|---|
| 26.0±0.8 | | 416, 652 | 1.16±0.33 (6/8) | 0.15±0.08 (2/3) | 0±0 (0/3) |
| 26.4 | | 460, 656 | 0.04±0.04 (1/8) | 0±0 (0/3) | 0±0 (0/3) |
| 27.0±0.4 | | 420, 464, 652 | 0.09±0.09 (1/8) | 0.09±0.09 (1/3) | 0±0 (0/3) |
| 27.3 | | 460, 648 | 0±0 (0/8) | 0.24±0.24 (1/3) | 0±0 (0/3) |
| **27.9±0.4** | **Allomer chl *b*** | **460, 648** | **15.24±1.59 (8/8)** | **3.75±0.91 (3/3)** | **0±0 (0/3)** |
| 27.9 | | 460, 648 | 0±0 (0/8) | 0.21±0.21 (1/3) | 0±0 (0/3) |
| 28.3 | | 464, 652 | 0.8±0.8 (1/8) | 0±0 (0/3) | 0±0 (0/3) |
| 29.0±0.3 | | 452, 636, 680 | 3.04±0.95 (6/8) | 0.42±0.23 (2/3) | 0±0 (0/3) |
| 29.4 | | 452, 636 | 0.32±0.32 (1/8) | 0±0 (0/3) | 0±0 (0/3) |
| 29.6±0.4 | | 452, 636 | 5.2±0.84 (8/8) | 0.98±0.39 (3/3) | 0±0 (0/3) |
| **30.5±0.5** | **Chl *b*** | **464, 648** | **1.01±0.2 (7/8)** | **15.64±2.59 (3/3)** | **0±0 (0/3)** |
| 31.6±0.4 | | 468, 652 | 0.48±0.14 (6/8) | 0±0 (0/3) | 0±0 (0/3) |
| 31.9 | | 432, 468, 668 | 0.05±0.05 (1/8) | 0±0 (0/3) | 0±0 (0/3) |
| **32.3±0.5** | **Allomer chl *b*** | **464, 648** | **0.37±0.16 (4/8)** | **4.12±0.75 (3/3)** | **0±0 (0/3)** |
| 33.0±0.4 | | 436, 672 | 0.07±0.05 (2/8) | 0±0 (0/3) | 0±0 (0/3) |
| **34.6±0.5** | **Allomer chl *a*** | **432, 664** | **12.08±1.64 (8/8)** | **4.29±1.68 (3/3)** | **0±0 (0/3)** |
| 35.6±0.4 | | colelution | 2.0±0.57 (8/8) | 0.47±0.24 (2/3) | 0±0 (0/3) |
| **36.6±0.4** | **Allomer chl *a*** | **420, 656** | **7.09±1.06 (8/8)** | **1.30±0.77 (3/3)** | **0±0 (0/3)** |
| **37.2±0.5** | **Chl *a*** | **432, 664** | **6.3±1.66 (7/8)** | **21.74±2.43 (3/3)** | **0±0 (0/3)** |
| **38.8±0.5** | **Allomer chl *a*** | **432, 668** | **3.19±0.43 (8/8)** | **6.06±0.40 (3/3)** | **0±0 (0/3)** |
| 40.8 | | 432, 668 | 0±0 (0/8) | 0.08±0.08 (1/3) | 0±0 (0/3) |
| 46.6 | | 468, 652 | 0.01±0.01 (1/8) | 0±0 (0/3) | 0±0 (0/3) |
| 48.2 | | 408, 664 | 0±0 (0/8) | 0.78±0.78 (1/3) | 0±0 (0/3) |
| 51.1 | | 468, 652 | 0.06±0.06 (1/8) | 0±0 (0/3) | 0±0 (0/3) |
| 52.5±0.5 | | 452, 480 | 3.71±0.45 (8/8) | 3.86±0.21 (3/3) | 0±0 (0/3) |
| 53.2 | | 444, 472 | 0.02±0.02 (1/8) | 0±0 (0/3) | 0±0 (0/3) |
| 53.7±0.3 | | 332, 348, 368, 444, 476 | 0.03±0.02 (2/8) | 0±0 (0/3) | 0±0 (0/3) |
| 54.6 | | 468, 648 | 0.05±0.05 (1/8) | 0±0 (0/3) | 0±0 (0/3) |

**Table S2: Chlorophyll *a* and *b* contents, and corresponding chl *b*: chl *a* ratios in endolithic *versus* free-living *Ostreobium* (clade P1).** Mean values ± Standard Error, SE; subcult.: replicate subculture.

| | **Strain** | **Chl *a* (µg/mg of organic extract)** | **Chl *b* (µg/mg of organic extract)** | **chl *a* : chl *b* ratio** | **chl *b* : chl *a* ratio** | **(chl *a* + its allomers) : (chl b + its allomers) ratio** | **(chl *b* + its allomers) : (chl a + its allomers) ratio** |
|---|---|---|---|---|---|---|---|
| **Free-living *Ostreobium* clade P1** | 05 | 0.84 | 0.19 | 4.5 | 0.22 | 1.42 | 0.7 |
| | 010 | 0 | 0.45 | 0 | / | 1.1 | 0.91 |
| | 018B subcult. n°1 | 26.8 | 2.92 | 9.16 | 0.11 | 2.91 | 0.34 |
| | 018B subcult. n°2 | 1.99 | 0.52 | 3.85 | 0.26 | 1.77 | 0.56 |
| | 019 subcult. n°1 | 0.77 | 0.2 | 3.89 | 0.26 | 1.47 | 0.68 |
| | 019 subcult. n°2 | 15.4 | 1.73 | 8.91 | 0.11 | 1.66 | 0.6 |
| | 018C subcult. n°1 | 4.26 | 0.69 | 6.21 | 0.16 | 1.55 | 0.64 |
| | 018C subcult. n°2 | 2.87 | 1.1 | 2.61 | 0.38 | 1.9 | 0.53 |
| **Mean ± SE (n=8)** | | **6.62±3.37** | **0.97±0.33** | **4.89±1.10** | **0.21±0.03** | **1.72±0.19** | **0.62±0.06** |
| **Endolithic *Ostreobium* clade P1** | 010 | 5.73 | 6.63 | 0.86 | 1.16 | 1.18 | 0.85 |
| | 018B | 17.6 | 9.53 | 1.85 | 0.54 | 1.46 | 0.68 |
| | 019 | 12.89 | 10.61 | 1.21 | 0.82 | 1.13 | 0.88 |
| **Mean ± SE (n=3)** | | **12.07±3.45** | **8.92±1.19** | **1.31±0.29** | **0.84±0.18** | **1.26±0.10** | **0.81±0.06** |

**Table S3: % level of fatty acids in *Ostreobium* strains, as endolithic or free-living filaments**. Data (Means±Standard Error, SE) are given for *Ostreobium* strains in clade P1 and in pooled clades P12/P14. Substrate control for endoliths is bleached coral carbonate skeleton. Data for the single endolithic strain representative of clade P14 are indicative and should be treated with caution because of lack of replication.

| Fatty acids | Free-living *Ostreobium* (% of total FA: mean±SE) | | Endolithic *Ostreobium* (% of total FA: mean±SE) | | |
|---|---|---|---|---|---|
| | clade P1 (n=8) | clade P12/P14 (n=5) | clade P1 (n=7) | clade P14 (n=1) | Control bleached skeleton (n=4) |
| **12:0** | 0.06±0.04 | 0.02±0.02 | 0.20±0.08 | 0 | 0.62±0.19 |
| **14:0** | 2.10±0.40 | 1.76±0.12 | 4.36±0.58 | 1.02 | 6.73±0.46 |
| **15:0iso** | 0.06±0.02 | 0.05±0.03 | 0 | 0 | 0 |
| **15:0anteiso** | 0.04±0.03 | 0.02±0.02 | 0 | 0 | 0 |
| **15:0** | 0.10±0.05 | 0.02±0.02 | 0.27±0.07 | 0 | 2.49±0.13 |
| **16:0iso** | 0.03±0.02 | 0 | 0 | 0 | 0 |
| **16:0** | 29.97±6.83 | 20.74±0.83 | 45.84±7.40 | 12.9 | 73.60±3.68 |
| **16:1ω9** | 0.30±0.07 | 0.25±0.08 | 0.42±0.15 | 0.39 | 0 |
| **16:1ω7** | 0.43±0.06 | 0.28±0.05 | 1.27±0.74 | 0.26 | 0 |
| **16:2ω6** | 0.87±0.10 | 0.61±0.11 | 2.73±1.11 | 0.62 | 0 |
| **16:3ω6** | 3.23±0.52 | 2.86±0.30 | 4.65±1.79 | 4.14 | 0 |
| **17:0** | 0.04±0.02 | 0 | 0.04±0.03 | 0.03 | 0 |
| **16:4ω3** | 0 | 0 | 0.04±0.04 | 0.39 | 0 |
| **18:0** | 2.27±0.30 | 1.11±0.26 | 7.29±1.67 | 2.24 | 16.56±3.82 |
| **18:1ω9** | 3.58±0.35 | 1.85±0.22 | 3.73±0.83 | 2.3 | 0 |
| **18:1ω7** | 5.72±0.42 | 4.48±0.19 | 3.21±0.87 | 3.29 | 0 |
| **18:2ω6** | 5.28±0.36 | 4.64±0.21 | 12.78±4.60 | 5.74 | 0 |
| **18:3ω6** | 2.64±0.49 | 2.16±0.27 | 4.09±1.45 | 3.28 | 0 |
| **19:1ω9** | 0.09±0.06 | 0 | 0 | 0 | 0 |
| **20:0** | 0 | 0.07±0.04 | 0.20±0.11 | 0 | 0 |
| **20:1ω9** | 0.05±0.02 | 0.02±0.02 | 0.06±0.06 | 0 | 0 |
| **20:2ω9** | 0.12±0.04 | 0.07±0.04 | 0.16±0.10 | 0.08 | 0 |
| **20:2ω6** | 0.14±0.07 | 0.30±0.12 | 0.03±0.02 | 0 | 0 |
| **20:3ω6** | 0.54±0.09 | 0.39±0.11 | 0.10±0.06 | 0.48 | 0 |
| **20:4ω6** | 41.47±8.42 | 57.25±0.45 | 7.84±2.69 | 60.31 | 0 |
| **20:5ω3** | 0.13±0.10 | 0.17±0.11 | 0.21±0.13 | 1.57 | 0 |
| **22:0** | 0.13±0.05 | 0.11±0.07 | 0.14±0.08 | 0.25 | 0 |
| **22:4ω6** | 0.30±0.11 | 0.23±0.14 | 0 | 0.66 | 0 |
| **22:5ω6** | 0.16±0.06 | 0.15±0.09 | 0.11±0.07 | 0 | 0 |
| **22:6ω3** | 0.20±0.13 | 0.39±0.25 | 0.18±0.14 | 0.17 | 0 |
| **24:0** | 0 | 0 | 0.05±0.05 | 0 | 0 |
| **ΣSFA** | 34.8±7.40 | 23.90±0.74 | 58.39±8.90 | 16.44 | 100±0 |
| **ΣMUFA** | 10.15±0.75 | 6.88±0.50 | 8.69±2.06 | 6.24 | 0 |
| **ΣPUFA** | 55.06±7.96 | 69.21±0.85 | 32.91±7.36 | 77.44 | 0 |
| **Σω3** | 0.32±0.21 | 0.56±0.36 | 0.42±0.25 | 2.13 | 0 |
| **Σω6** | 54.62±7.82 | 68.58±0.94 | 32.33±7.12 | 75.23 | 0 |

**Table S4: C/N ratios of individual *Ostreobium* strains.** Endolithic filaments were analyzed after decalcification with formic acid 5% (decalcified endolithic *Ostreobium*). Control substrate of endoliths was skeletal organic matrix. NA: non available. Mean values ± Standard Deviation, SD. Subcult.: replicate subculture, techn. replicates: technical replicates.

| | Strain | *rbc*L clade | C/N ratio | acid-treated C/N ratio |
|---|---|---|---|---|
| **Free-living *Ostreobium*** | 05 subcult. n°1 | P1 | 12.4 | NA |
| | 05 subcult. n°2 | P1 | 13.5 | NA |
| | 010 subcult. n°1 (techn. replicate n°1) | P1 | 17.3 | NA |
| | 010 subcult. n°1 (techn. replicate n°2) | P1 | 17.7 | NA |
| | 010 subcult. n°1 (techn. replicate n°3) | P1 | 17.9 | NA |
| | 018B | P1 | 20.5 | NA |
| | 019 | P1 | 17.0 | NA |
| | 022 subcult. n°1 | P1 | 14.5 | NA |
| | 022 subcult. n°2 | P1 | 16.2 | NA |
| | 018A subcult. n°1 | P12 | 14.1 | NA |
| | 018A subcult. n°2 | P12 | 13.6 | NA |
| | 06 subcult. n°1 | P14 | 16.4 | NA |
| | 06 subcult. n°2 | P14 | 13.2 | NA |
| **Mean ± SD (all strains, n=13)** | | | **15.7±2.4** | NA |
| **Mean ± SD (clade P1, n=9)** | | | **16.3±2.5** | NA |
| **Mean ± SD (clade P12/P14, n=4)** | | | **14.3±1.4** | NA |
| **Decalcified endolithic *Ostreobium*** | 05 subcult. n°1 | P1 | | 12.2 |
| | 05 subcult. n°2 | P1 | | 12.6 |
| | 010 subcult. n°1 | P1 | | 13.5 |
| | 022 subcult. n°1 | P1 | | 12.8 |
| | 022 subcult. n°2 | P1 | | 11.9 |
| | 018A subcult. n°1 | P12 | | 11.9 |
| | 018A subcult. n°2 | P12 | | 12.3 |
| | 06 subcult. n°1 | P14 | | 11.5 |
| | 06 subcult. n°2 | P14 | | 12.0 |
| **Mean ± SD (all strains, n=9)** | | | | **12.3±0.6** |
| **Mean ± SD (clade P1, n=5)** | | | | **12.6±0.6** |
| **Mean ± SD (clade P12/P14, n=4)** | | | | **11.9±0.3** |
| **Skeletal organic matrix** | / | / | | 12.3 |
| | / | / | | 11.0 |
| | / | / | | 11.3 |
| | / | / | | 9.7 |
| **Mean ± SD (n=4)** | | | | **11.1±1.1** |

**Table S5 : δ$^{13}$C and δ$^{15}$N values of individual *Ostreobium* strains.** Endolithic filaments were analyzed within their carbonate substrate (non decalcified endolithic *Ostreobium*) or after decalcification with formic acid 5% (decalcified endolithic *Ostreobium*). δ$^{13}$C values corrected by acid treatment (+6‰) for decalcified endolithic *Ostreobium* are also provided in italics. Control substrate of endoliths was either bleached coral carbonate skeleton (composed of $CaCO_3$ and organic matrix) or skeletal organic matrix. ND: δ$^{15}$N was below EA-IRMS detection limit. NA: non available. Mean values ± Standard Deviation, SD. Subcult.: replicate subculture, techn. replicates: technical replicates.

| | **Strain** | ***rbc*L clade** | **δ$^{13}$C (‰)** | **acid-treated δ$^{13}$C (‰)** | **δ$^{15}$N (‰)** | **acid -treated δ$^{15}$N (‰)** |
|---|---|---|---|---|---|---|
| **Free-living *Ostreobium*** | 05 subcult. n°1 | P1 | -16.2 | NA | 5.0 | NA |
| | 05 subcult. n°2 | P1 | -16.9 | NA | 4.8 | NA |
| | 010 subcult. n°1 (techn. replicate n°1) | P1 | -18.6 | NA | 2.6 | NA |
| | 010 subcult. n°1 (techn. replicate n°2) | P1 | -21.7 | NA | 3.7 | NA |
| | 010 subcult. n°1 (techn. replicate n°3) | P1 | -21.7 | NA | 3.2 | NA |
| | 010 subcult. n°2 (techn. replicate n°1) | P1 | -23.4 | -28.2 | 1.8 | 1.1 |
| | 010 subcult. n°2 (techn. replicate n°2) | P1 | -21.1 | -28.6 | 2.3 | 1.9 |
| | 010 subcult. n°2 (techn. replicate n°3) | P1 | -18.2 | -23.8 | 2.8 | 3.5 |
| | 018B | P1 | -18.5 | NA | 2.8 | NA |
| | 019 | P1 | -20.3 | NA | 1.4 | NA |
| | 022 subcult. n°1 | P1 | -17.4 | NA | 5.2 | NA |
| | 022 subcult. n°2 | P1 | -16.4 | NA | 5.4 | NA |
| | 018A subcult. n°1 | P12 | -17.2 | NA | 7.4 | NA |
| | 018A subcult. n°2 | P12 | -19.8 | NA | 9.0 | NA |
| | 06 subcult. n°1 | P14 | -14.7 | NA | 5.3 | NA |
| | 06 subcult. n°2 | P14 | -14.9 | NA | 6.3 | NA |
| **Mean ± SD (all strains, n=16)** | | | **-18.6±2.6** | NA | **4.3±2.1** | NA |
| **Mean ± SD (clade P1, n=12)** | | | **-19.2±2.4** | NA | **3.4±1.4** | NA |
| **Mean ± SD (clade P12/P14, n=4)** | | | **-16.7±2.4** | NA | **7.0±1.6** | NA |
| **Non decalcified endolithic *Ostreobium*** | 010 subcult. n°1 (techn. replicate n°1) | P1 | -10.9 | | NA | |
| | 010 subcult. n°1 (techn. replicate n°2) | P1 | -12.2 | | 4.6 | |
| | 010 subcult. n°1 (techn. replicate n°3) | P1 | -10.4 | | 3.6 | |
| | 010 subcult. n°1 (techn. replicate n°4) | P1 | -11.1 | | 4.5 | |
| **Mean ± SD (clade P1, n=4)** | | | **-11.2±0.8** | | **4.2±0.6** | |

| | | | | | | |
|---|---|---|---|---|---|---|
| **Decalcified endolithic *Ostreobium*** | 05 subcult. n°1 | P1 | *-20.7* | -26.7 | | 17.9 |
| | 05 subcult. n°2 | P1 | *-21.2* | -27.2 | | 13.0 |
| | 010 subcult. n°1 | P1 | *-15* | -21.0 | | 4.7 |
| | 010 subcult. n°2 (techn. replicate n°1) | P1 | *-14.3* | -20.3 | | 4.3 |
| | 010 subcult. n°2 (techn. replicate n°2) | P1 | *-13.2* | -19.2 | | 5.0 |
| | 010 subcult. n°2 (techn. replicate n°3) | P1 | *-13.2* | -19.2 | | 5.4 |
| | 010 subcult. n°2 (techn. replicate n°4) | P1 | *-14.3* | -20.3 | | 5.0 |
| | 022 subcult. n°1 | P1 | *-19* | -25.0 | | 11.2 |
| | 022 subcult. n°2 | P1 | *-17.8* | -23.8 | | 16.8 |
| | 018A subcult. n°1 | P12 | *-22.7* | -28.7 | | 16.2 |
| | 018A subcult. n°2 | P12 | *-22.6* | -28.6 | | 10.0 |
| | 06 subcult. n°1 | P14 | *-16* | -22.0 | | 17.1 |
| | 06 subcult. n°2 | P14 | *-20.9* | -26.9 | | 12.1 |
| **Mean ± SD (all strains, n=13)** | | | ***-17.8±3.6*** | **-23.8±3.6** | | **10.7±5.3** |
| **Mean ± SD (clade P1, n=9)** | | | ***-16.5±3.2*** | **-22.5±3.2** | | **9.3±5.6** |
| **Mean ± SD (clade P12/P14, n=4)** | | | ***-20.5±3.1*** | **-26.5±3.1** | | **13.9±3.4** |
| **Control coral skeleton** | / | / | -13.9 | | ND | |
| | / | / | -13.9 | | ND | |
| | / | / | -13.9 | | ND | |
| **Mean ± SD (n=3)** | | | **-13.9±0.03** | | ND | |
| **Skeletal organic matrix** | / | / | | -30.1 | | 5.5 |
| | / | / | | -28.8 | | 5.1 |
| | / | / | | -30.9 | | 5.0 |
| | / | / | | -27.9 | | 6.2 |
| **Mean ± SD (n=4)** | | | | **-29.4±1.3** | | **5.5±0.5** |

**Table S6: Turnover rates of Carbon (C) and Nitrogen (N) in endolithic *versus* free-living *Ostreobium*, in light and dark conditions.** Turnover values expressed in % were calculated for a 12h daytime period, extrapolated from the isotope ratio values (*enriched; *ctl*: normal abundance) measured at the end of the 8h labeling pulse with $^{13}C$-bicarbonate and $^{15}N$-nitrate, with the equation from Kopp *et al.* (2015): $[(((^{13}C/^{12}C)^{*}-(^{13}C/^{12}C)^{ctl}) \times 12) / ((1+(^{13}C/^{12}C)^{ctl}) \times 8)] \times 100$.
For endolithic *Ostreobium*, isotopic ratio values were corrected by subtracting the average values of 3 replicates control non-colonized labeled skeletons (corresponding to non-specific absorption of the isotopic label on the skeletal matrix). techn. replicates: technical replicates.

Kopp, C., Domart-Coulon, I., Humbell, B., Escrig, S., Hignette, M., and Meibom, A. (2015) Subcellular investigation of photosynthesis-driven carbon and nitrogen assimilation and utilization in the symbiotic reef coral *Pocillopora damicornis*. *mBIO* 6: e02299-14.

| | | | Light conditions ☼ | | Dark conditions ☾ | |
|---|---|---|---|---|---|---|
| | **Strain** | ***rbc*L clade** | **%Turnover of C (/12h daytime)** | **% Turnover of N (/12h daytime)** | **% Turnover of C (/12h daytime)** | **% Turnover of N (/12h daytime)** |
| **Free-living *Ostreobium*** | 022 | P1 | 1.45 | 1.31 | 0.04 | 0.86 |
| | 05 | P1 | 3.43 | 0.96 | 0.04 | 0.83 |
| | 010 – techn. replicates n°1 | P1 | 0.94 | 0.21 | NA | NA |
| | 010 – techn. replicates 2 | P1 | 0.86 | 0.18 | NA | NA |
| | 010 – techn. replicates 3 | P1 | 0.90 | 0.23 | NA | NA |
| | 018A | P12 | 4.65 | 1.36 | 0.08 | 0.83 |
| | 06 | P14 | 3.63 | 1.55 | 0.04 | 0.92 |
| **Mean ± SD (all strains, n=7)** | | | **2.27±1.59** | **0.83±0.61** | **0.05±0.02** | **0.86±0.04** |
| **Mean ± SD (clade P1, n=5)** | | | **1.52±1.10** | **0.58±0.52** | **0.04±0.01** | **0.84±0.02** |
| **Mean ± SD (clade P12/P14, n=2)** | | | **4.14±0.73** | **1.45±0.13** | **0.06±0.02** | **0.87±0.07** |
| **Endolithic *Ostreobium*** | 022 | P1 | 1.94 | 0.80 | 0.01 | 0.13 |
| | 05 | P1 | 2.42 | 0.91 | 0.04 | 0.62 |
| | 010 | P1 | 1.28 | 0.75 | NA | NA |
| | 018A | P12 | 0.48 | 0.24 | 0.03 | 0.21 |
| | 06 | P14 | 1.82 | 0.84 | 0.02 | 0.30 |
| **Mean ± SD (all strains, n=5)** | | | **1.59±0.74** | **0.71±0.27** | **0.02±0.02** | **0.32±0.21** |
| **Mean ± SD (clade P1, n=3)** | | | **1.88±0.57** | **0.82±0.08** | **0.02±0.02** | **0.38±0.34** |
| **Mean ± SD (clade P12/P14, n=2)** | | | **1.15±0.95** | **0.54±0.43** | **0.03±0.01** | **0.26±0.07** |
